# AllMetal3D: joint prediction of localization, identity and coordination geometry of common metal ions in proteins

**DOI:** 10.1101/2025.02.05.636627

**Authors:** Simon L. Dürr, Ursula Rothlisberger

## Abstract

Nature uses a variety of metal ions as cofactors for specific tasks. The three main classes of metals in biological systems are alkali ions (e.g. sodium and potassium), alkaline earth ions (e.g. magnesium and calcium) as well as transition metal ions (e.g. zinc or iron and many others). To date no selective predictor to localize each of these metals in a given protein structure exists. Current methods require either preselecting a specific ion (e.g MIB2 (***Lu et al., 2022***), ESMBind (***Dai et al., 2024***) or AlphaFold3 (***Abramson et al., 2024***) or a location (MIC ***Shub et al. (2024)***). In this work, we describe an extension of the recently introduced Metal3D framework (***Dürr et al., 2023***) that was originally trained on zinc sites to predict the location of all biologically relevant classes of metal ions as well as to classify their coordination geometry. Our model is the first of its kind and even outperforms Metal3D for the prediction of the location of Zn^2+^ and generalizes well to the other metals in terms of location prediction as well as identity classification. Comparing the model to several other available tools such as MetalSiteHunter, AlphaFold3, MIC, MIB2 and MetalHawk highlights important shortcomings in these tools with respect to data bias, prediction of negative sites and selectivity. However, concerning coordination geometry prediction, similar to other work, we find that our method cannot accurately make correct classifications beyond the most common classes in natural proteins i.e. tetrahedral and octahedral arrangements. AllMetal3D is available as ChimeraX extension, standalone web app as well as python package: https://github.com/lcbc-epfl/allmetal3d

## Introduction

Metal ions are essential for biology and are involved in enzyme catalysis, signal transduction and in regulating proteins (***Andreini et al., 2008***; ***Waldron and Robinson, 2009***). The biologically most relevant metal ions are alkali, alkaline-earth and transition metal ions. While alkali ions are mainly involved in signal transduction processes and bind mostly transiently to proteins (***Sigel et al., 2016***), alkaline-earth ions fulfill dual roles in signal transduction and as protein co-factors (***Yang, 2011***; ***Sigel and Sigel, 2019***). Calcium is less used for catalysis whereas magnesium is essential for nucleobase chemistry (***Yang, 2011***). Transition metal ions on the other hand are mostly involved in enzyme catalysis with zinc e.g. being typically used as Lewis acid, iron for redox reactions and copper for oxidative phosphorylation (***Sigel and Sigel, 2019***; ***Crans and Kostenkova, 2020***). We recently introduced the Metal3D framework for predicting metal ion locations and trained it on zinc finding that it could also predict other transition metal locations well (***Dürr et al., 2023***). However, Metal3D does not perform well for alkaline-earth or alkali ions and also cannot differentiate between different metal ions (***Dürr et al., 2023***). In this work, we present a new pipeline, AllMetal3D, an extension to the Metal3D framework trained on different metal ion binding sites to create a method, that selectively predicts the location, identity and geometry for all of the biologically most relevant metal ions, i.e. alkali, alkaline-earth- and transition metals.

### Selective location prediction

To the best of our knowledge, selective blind prediction of metal ion location and identity in a single computational pipeline for all biologically relevant metals has not been demonstrated yet. Some tools like MIB2 (***Lu et al., 2022***) or ESMBind (***Dai et al., 2024***) output predictions for multiple transition metal ions (ESMBind) or transition metals and alkaline-earth ions (MIB2). However, both tools are not designed to predict selectivity. MIB2 uses template similarity to existing metal binding sites in its database, which should impart it with some selectivity, however template similarity is not a direct proxy for selectivity especially since for some metal ions, there are only few structural templates available. PinMyMetal (***Zheng et al., 2024***) predicts location and selectivity for transition metal ions. Other tools using structural input are designed for either location prediction such as BioMetAll (***Sánchez-Aparicio et al., 2021***), binding site prediction such as MasterOfMetals (***Laveglia et al., 2023***) or identity prediction such as MetalSiteHunter (***Mohamadi et al., 2022***) and MIC (***Shub et al., 2024***). Existing 2D sequence approaches exhibit many known biases (***Walsh et al., 2016***) and only annotate putative binding residues for different metal ions. Cofolding methods like AlphaFold3 (***Abramson et al., 2024***), Chai-1 (***Chai Discovery et al., 2024***), RoseTTAFold-AllAtom (***Krishna et al., 2024***) and Boltz-1 (***Wohlwend et al., 2024***) require input of the specific metal ion and cannot directly be used for selectivity prediction (***Dürr and Rothlisberger, 2024***).

Several other location predictors like PinMyMetal (***Zheng et al., 2024***), ESMBind (***Dai et al., 2024***) or MasterOfMetals (***Laveglia et al., 2023***) have appeared since Metal3D was released but they also are trained only on transition metals (ESMBind, PinMyMetal) or Zn^2+^only. Sidechain based predictions such as used by PinMyMetal preclude prediction of backbone coordinated alkali or alkaline-earth ions because those can be bound to all backbone carbonyls independent of the side chain.

### Identity prediction

***Mohamadi et al. (2022)*** developed MetalSiteHunter, a 3DCNN model for classifying metal ion identities given a location. The model uses similar input channels as Metal3D removing the hydrophobic and aromatic channels that were not deemed important based on feature selection. MetalSiteHunter uses a balanced dataset of 1260 examples each for the metal ions Zn^2+^, Fe^3+^, Mg^2+^, Mn^2+^, Na^+^ and non-metal sites as a negative training set. The non-metal dataset is created by sampling pockets in proteins located using fpocket (***Le Guilloux et al., 2009***). The model is trained using 5-fold cross validation. Each site is sampled with the voxel grid centered on the average position of all C_*α*_ atoms of metal binding residues of a given metal site and not on the specific location of the ion itself.

Metric Ion Classification (MIC) (***Shub et al., 2024***) uses interaction fingerprints to classify positions in proteins as metal ions, halide ions or water sites. Two models are available: One trained on Mg^2+^, Zn^2+^, Ca^2+^, Na^+^, Cl^-^ and water, the other MIC_extended_ trained on Mg^2+^, Zn^2+^, Ca^2+^, Na^+^, Cl^−^, K^+^, Fe^3+^, Cl^-^ and water. Both models were shown to perform well on a set of manually curated examples. However, waters close to metal binding sites are often misidentified as metal ions.

CheckMyBlob (***Kowiel et al., 2018***; ***Brzezinski et al., 2021***) uses the electron density map to classify density peaks. It does not need input locations for the prediction like MIC or MetalSiteHunter and classifies the density peaks directly. CheckMyBlob classifies 219 different ligands and groups them in categories such as Zn^2+^-like or Ca^2+^-like. The final model is an ensemble of a k-nearest neighbors (k-NN), random forest (RF), and gradient boosting machine (GBM) with a GBM aggregating the final predictions. 60 features are used to make predictions, which were chosen out of a set of 382 features through recursive feature elimination (***Kowiel et al., 2018***). These features include e.g. the resolution of the crystal structure, the number of adjacent oxygen or nitrogen atoms to the blob, features computed on the electron density such as the gradient between different contour levels or Zernike moment invariants (***Novotni and Klein, 2004***). A webserver is available (***Brzezinski et al., 2021***) but no code or API is provided to run single predictions.

### Coordination geometry prediction

Many metal ions can exist in different oxidation states and some, such as calcium or zinc, can bind with different coordination numbers and geometries (***Bazayeva et al., 2024***). In fact, changes in the number of coordinating atoms or in the coordination geometry are common for enzymatically catalyzed reactions and are often used to drive the catalytic cycle (***Moura et al., 2008***; ***Dürr et al., 2021***).

For the automated assignment of coordination number and geometry, multiple methods and models exists. To construct the MetalPDB (***Putignano et al., 2018***) database, FindGeo (***Andreini et al., 2012***) was used. FindGeo, assigns coordination geometry via superposition with geometries of ideal coordination polyhedra. A similar method is CheckMyMetal (***Zheng et al., 2017***), which verifies geometry by comparing to ideal polyhedra via a metric termed gRMSD. Another approach, mainly intended for inorganic crystals, is ChemEnv in pymatgen (***Waroquiers et al., 2020***). ChemEnv uses Voronoi analysis to assign nearest neighbors of the target metal ion and compares the polyhedra formed by the nearest neighbor atoms to assign a coordination geometry.

Only one recently published deep learning based method exists to assign coordination geometries. MetalHawk (***Sgueglia et al., 2024***) is a fully connected neural network using the six nearest atoms to the metal ion to predict the coordination geometry. The model is fed a distance matrix and an angle matrix and can predict 7 different coordination environments. Two different datasets are used to train the model: the Cambridge structure database (CSD) and the protein data bank (PDB). The model was shown to work better using the CSD dataset, which the authors attribute to data sparsity in the PDB for less populated classes other than tetrahedral or octahedral sites. The model is trained using only transition metal ions.

In this work, we integrate multiple approaches to extend the initial Metal3D model to a broad range of biologically relevant metal ions. We explore a model that selectively predicts localization for different metal ions in different channels and a simpler model that predicts a general metal density and then classifies the probability peaks with respect to identity, geometry and presence of vacant sites.

## Results

We set out to train a single unified pipeline that given a single protein structure can predict the location, identity and likely coordination geometry of biologically relevant metal ions. The basis for this pipeline is the Metal3D framework that makes use of fully convolutional 3DCNNs (***Dürr et al., 2023***). We split training of the models into two different subtasks: location prediction using the Metal3D framework and classification of the predicted locations given a fingerprint of the environment around the predicted site (Figure 1).

**Figure 1.**
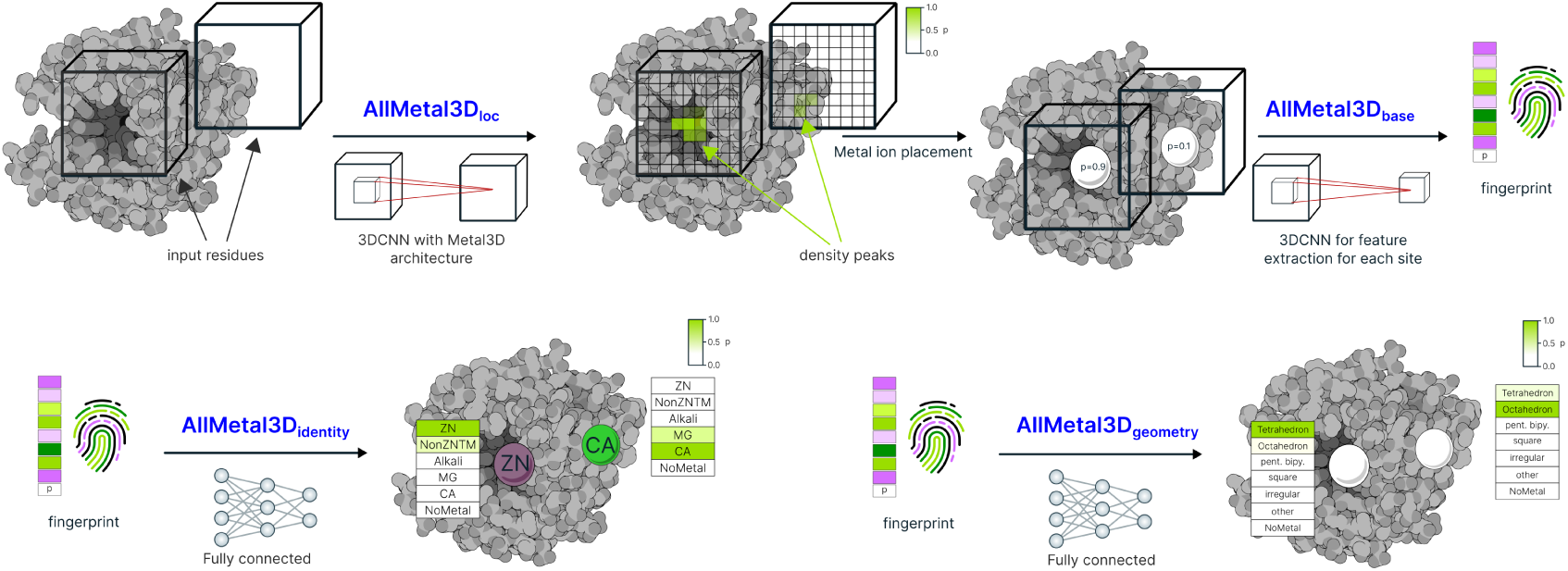
Architecture of the AllMetal3D pipeline composed of AllMetal3D_*loc*_ with modified dataset compared to Metal3D and AllMetal3D_base_ learning a fingerprint for each predicted location and two small fully connected neural networks: AllMetal3D_identity_ and AllMetal3D_geometry_.

### Location prediction

First, we trained a new version of Metal3D by modifying only the input dataset, i.e. extending the zinc only based training set to other metalloproteins. This model, which we term AllMetal3D_loc_ predicts all metals in a single output channel. This means that the output density map now encodes the probability of a general metal being bound at a specific location.

We employed a similar set of structures used for testing selectivity of Metal3D (***Dürr et al., 2023***) to evaluate whether AllMetal3D_loc_ can predict the extended range of metal ions with higher precision and recall. Interestingly, overall the probability for predicted sites is a bit lower for AllMetal3D than for Metal3D (Figure S8). Based on the lowered probability for metal site prediction, we used

AllMetal3D_loc_ at a probability threshold of p=0.65 and Metal3D at the cutoff previously determined to work best for physiological sites of p=0.75 (***Dürr et al., 2023***). We also empirically determined that predicted metal coordinates are more precise when only including probability densities above p=0.25 and not p=0.15 as for Metal3D.

Figure 2 shows that the performance of AllMetal3D_loc_ improves both in terms of precision and recall for location prediction for all metals where performance has been suboptimal for Metal3D. Especially for Ca^2+^, the performance is much improved. Performance improvements for Mg^2+^ are less pronounced with an increase in recall from 0.2 to 0.43 for Metal3D. The performance for K^+^ and Na^+^ is now almost equal. For transition metals, performance does only slightly change with zinc still being the metal with highest combination of precision and recall. Like for Metal3D performance is best on metals coordinated by at least 3 unique protein residues (Figure 2) with pronounced performance differences with respect to 2+ coordinated metals for all metal ions. The performance on Zn^2+^ is better across the board for AllMetal3D than Metal3D despite being trained also on other metal ions (Figure S1). Among the other metals, performance is best for Ca^2+^ and Mn^2+^. Recall for Co^2+^ is worse than for other transition metals whereas precision at high probability threshold is higher.

**Figure 2.**
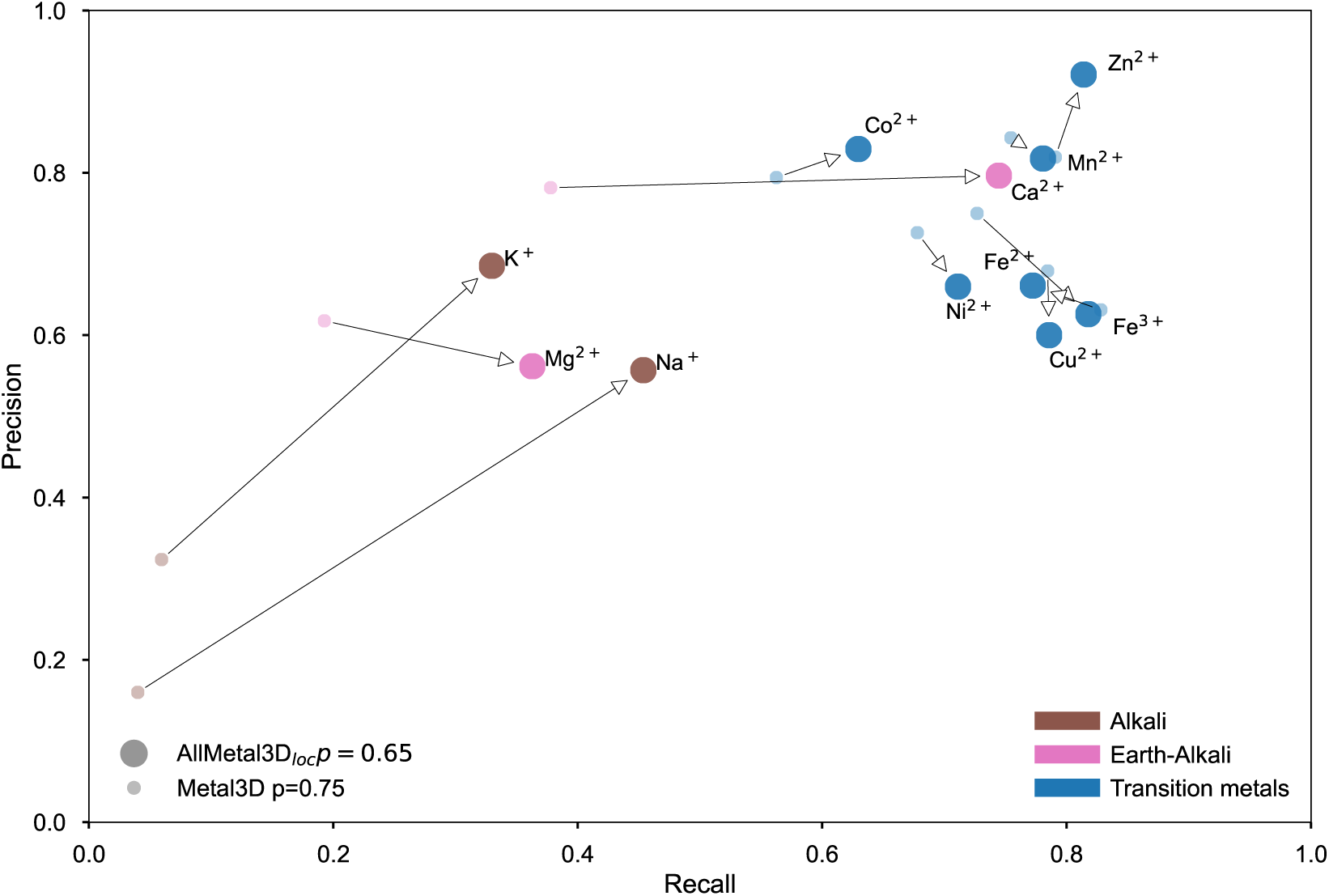
Precision and Recall for physiological binding sites (with 3+ unique coordinating protein residues) for all biologically important alkali, alkaline-earth and transition metal ions with p=0.75 for Metal3D and p=0.65 for AllMetal3D_loc_. Arrows indicate the performance difference between Metal3D and AllMetal3D_loc_, which is evaluated on location only and does not predict the identity.

Metal ions with high precision and recall are predicted with high spatial precision. Predictions are most spatially precise with a median of 0.5 Å for Ca^2+^, Ni^2+^, Cu^2+^ and Zn^2+^ (Figure S2). The predictions for Fe^3+^ and Mn^2+^ are less spatially precise with two peaks around 0.5 Å and 1.2 Å. Predictions for Mg^2+^ and Na^+^ and K^+^ are equally good compared to Fe^3+^ and Mn^2+^ with median of 0.6 Å.

Similarly, the average probability for the correctly identified predicted sites is lower for Mg^2+^ and Na^+^ and K^+^ than for the other ions (Figure S8). Ca^2+^ is the most confidently predicted metal ion with a median probability of 0.98 and mean absolute deviation of 0.94 ± 0.08 Å. Among the transition metals, the model is confident for all of them but less in comparison to Metal3D on the same metals (***Dürr et al., 2023***). For example for Zn^2+^ the mean probability of AllMetal3D is p=0.90 ± 0.07 compared to p=0.97 ± 0.05 in the case of Metal3D (Table S3).

### Classifying negative sites

AllMetal3D performs two different types of classification: AllMetal3D_loc_ performs binary classification for each point whether it contains a metal ion or not. AllMetal3D_identity_ and AllMetal3D_geometry_ perform classification per metal ion given an environment around the metal. It is important for classifiers to be robust to negative samples.

Therefore, we also analyzed how non-metal binding pockets in proteins are classified. We compared AllMetal3D_*loc*_, MIB2 (***Lu et al., 2022***) and MetalSiteHunter (***Mohamadi et al., 2022***) on this task. We sampled non-metal binding pockets using fpocket (***Le Guilloux et al., 2009***) analyzing the center of the deepest pocket in a given protein. MetalSiteHunter (***Mohamadi et al., 2022***) uses fpocket in the same way to create negative samples for training their model.

Non-metalloproteins were input into AllMetal3D_loc_ and we then checked whether the model does not predict density in the deepest pocket of a non-metalloprotein using a cutoff of 5 Å to the center of the pocket. MIB2 makes predictions on whole proteins and we checked whether any metals are predicted inside the deepest pocket of a non-metalloprotein using a t-score cutoff of t=2.5. MetalSiteHunter classifies the environment instead of providing location predictions, so here we used the center of all atoms lining the pocket from the fpocket output (***Le Guilloux et al., 2009***) and checked whether the model predicts the NoMetal label. We evaluated AllMetal3D against MetalSiteHunter or MIB2 on the same sets of 528, respectively 113 structures. The set of 113 structures is contained in the set of 528 structures and is smaller due to computational constraints for submission to the MIB2 webserver.

For the small dataset, we found that comparing AllMetal3D_loc_ and MIB2, non-metal pockets were mostly consistently predicted as non-metal binding (Figure 3) with 17 false positives for MIB2 and none for AllMetal3D_loc_ above p=0.5. Out of the false predictions for MIB2, only one overlapped with the AllMetal3D_loc_ predictions with 0.3<p<0.5. The average t-score for false predictions of MIB2 is 4.02 ±1.15 with an average distance of the predicted ion position to the center of the pocket of 3.86 ± 0.89 Å. On a larger set of 528 non-metalloproteins and their largest pocket, MetalSiteHunter predicted only 183 of the expected 528 non-metal pockets as non-metal binding (Figure 3). The incorrectly predicted non-metal pockets were often classified as Mg^2+^ or Na^+^. Zn^2+^ is predicted in 2 cases. In contrast, AllMetal3D_loc_ just predicted 2 pockets with probability higher than 0.5 in this set.

**Figure 3.**
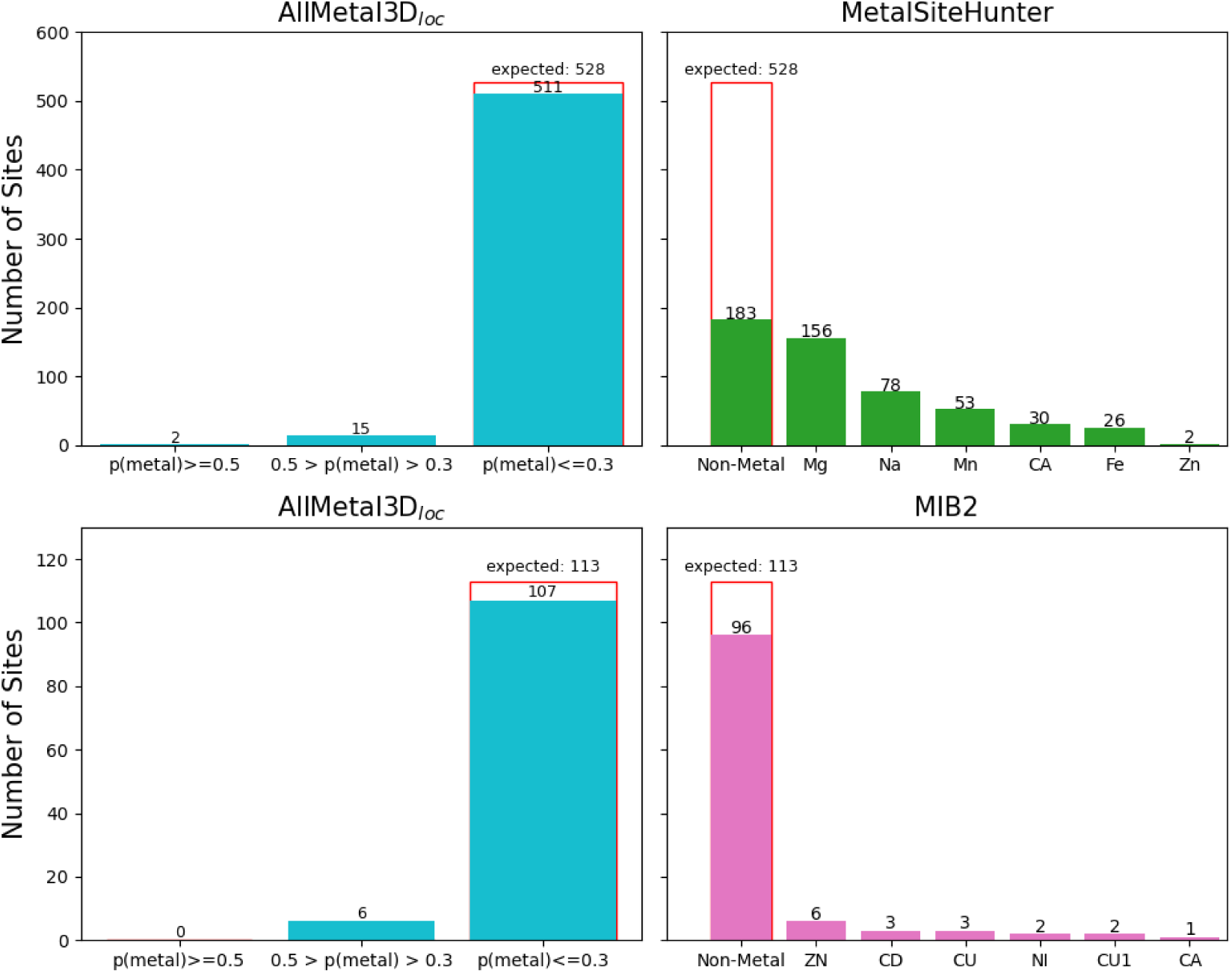
Performance of AllMetal3D_loc_, MetalSiteHunter and MIB2 for negative sites. Comparison is done on the largest non-metal pocket for two sets of 528 for MetalSiteHunter, respectively 113 non-metalloproteins for MIB2.

### Identity prediction

We ran the AllMetal3D_loc_ model on each structure used to construct the training and test sets for identity prediction. The ground truth labels were then assigned by comparing the predicted density peaks from AllMetal3D_loc_ with experimental sites. For each predicted location with probability above 0.25, we stored the coordinate along with the label of the closest metal in the input structure in the dataset used to train AllMetal3D_identity_. While we ran predictions with AllMetal3D_loc_ for generating the dataset for AllMetal3D_identity_ only on residues close to an experimental metal binding sites there can still be false positives contained in these predictions i.e. sites that are not within 5 Å of a metal in the crystal structure. For all false positive predictions we stored the label `NoMetal`.

We first trained a model on the full dataset containing 29844 entries for training and 6698 for testing using 3 fold rotational data augmentation with random uniform rotation. We did not correct for dataset imbalance between the different metal ion classes. This model performs well to classify Zn^2+^, Ca^2+^, and Mg^2+^ (Figure 4). Zn^2+^ and Ca^2+^ are the majority classes in the dataset. The accuracy of this model is 46% and a top-3 accuracy on the test set of 80% is reached. However, the less common ions in the test set are almost exclusively assigned to the most similar majority class. No K^+^, Fe^3+^ and Ni^2+^ sites are predicted. Most of the true Fe^3+^ and Ni^2+^ binding sites are assigned as Zn^2+^. Cu^2+^ is partly assigned to Zn^2+^ partly correct as Cu^2+^. Mn^2+^ is assigned as Zn^2+^, Ca^2+^ or Mg^2+^. Since such predictions are of little utility and might even mislead users to model the wrong ion instead of the true but rarer ion, we opted to combine categories and create a more balanced dataset of labels and retrain the model on this more balanced set of examples (see Figure S10 for statistics).

**Figure 4.**
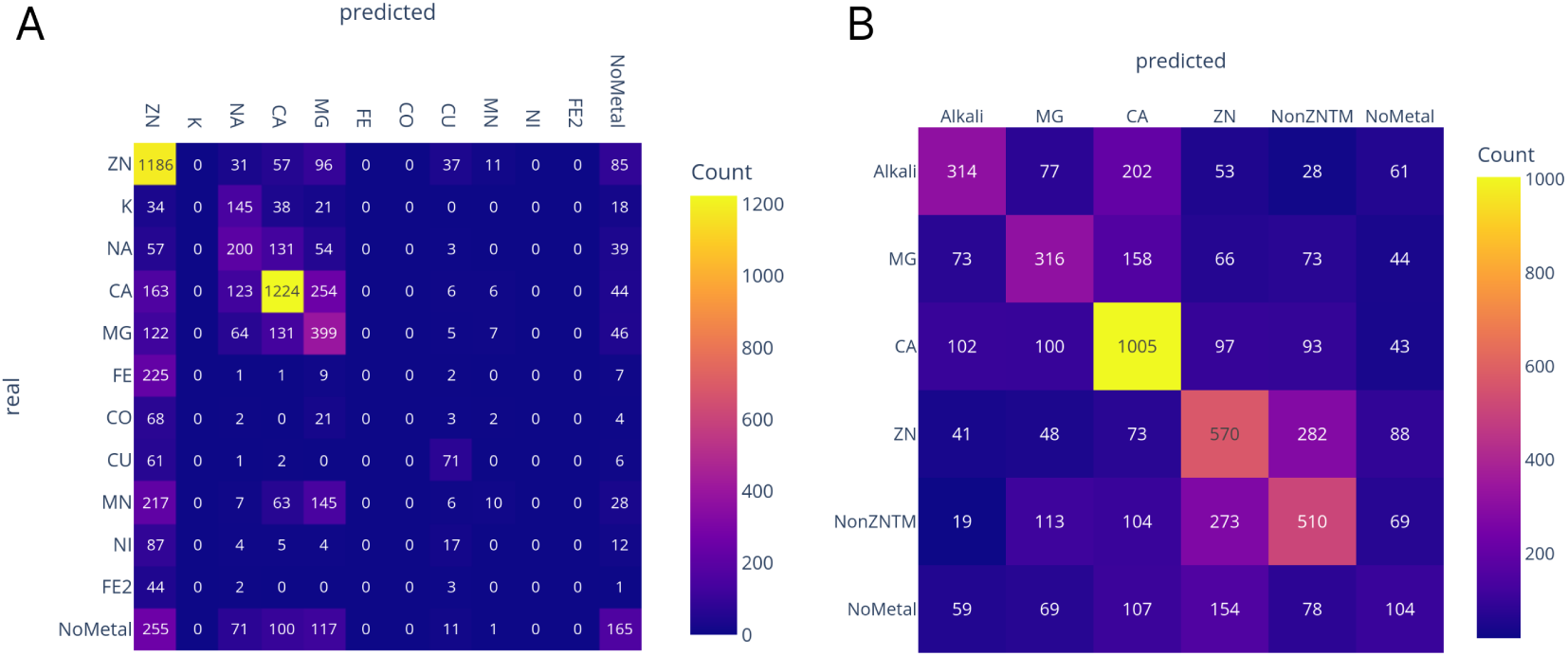
AllMetal3D_identity_ confusion matrix for test proteins when trained on A individual ion classes and B on merged ion classes.

The model trained on the reduced class set performs better but not perfectly (Figure 4B). However, the model makes sensible misclassifications where the main categories that are confused are Zn^2+^ with non-zinc transition metals as well as Ca^2+^ with Mg^2+^ and alkali ions. While for Zn^2+^ 51.7 % of predictions are correct with an additional 25.6% misclassified as non-zinc transition metals, for non-zinc transition metals 46.9 % are classified correctly and 25% are classified as Zn^2+^. Some of the real metal binding sites are also predicted as NoMetal. Most of the true NoMetal cases are predicted as Ca^2+^ or Zn^2+^ with just 104 out of 571 assigned correctly.

***Mohamadi et al. (2022)*** report good results on classification of Zn^2+^, K^+^, Na^+^, Mg^2+^, Ca^2+^, and non-metal sites for MetalSiteHunter. We constructed a subset of metal ions including false positive location predictions from the AllMetal3D test and used the MetalSiteHunter webserver to classify the center of all residues that coordinate a prediction from AllMetal3D_*loc*_. For this set, we included all predictions above p>0.1. On the resulting 2960 test metal sites, performance of MetalSiteHunter differs from the published performance of the authors (Figure S3). While Zn^2+^ and Ca^2+^ are confidently correctly assigned by MetalSiteHunter, the model has no selectivity for prediction of Mn^2+^ and seldomly predicts the false positive locations as non-metal binding inline with the low selectivity for non-metal pockets (Figure 3). Fe^3+^ is predicted mostly as Mn^2+^, Na^+^ as Mg^2+^ or Ca^2+^, and Mg^2+^ as Ca^2+^. Interestingly, however many of the false positive locations from AllMetal3D are predicted as Mg^2+^ consistent with the results on the set of negative examples (Figure 3).

MIB2 performs template search given a protein and list of metals to search for (***Lu et al., 2022***). MIB2 was not designed to selectively identify metals but we reasoned that higher template similarity could be a good proxy for selectivity prediction and therefore compared AllMetal3D_identity_ and MIB2 on a set of 1520 structures (Figure S4). In this set, we excluded alkali ions since MIB2 does not offer predictions for these ions. In view of the fact that MIB2 was not specifically designed to selectively predict metal ion identity, we also checked whether the correct metal ion was among the top-3 predictions with the highest template score. It turns out that while MIB2 does reasonably well in identifying Ca^2+^ and Zn^2+^ sites, it clearly misassigns many of the other transition metals, in particular a significant number of Zn^2+^ sites are predicted as Cu^2+^ or Fe^3+^. Surprisingly, even magnesium and calcium sites are frequently mispredicted as Fe^3+^, Zn^2+^, Co^2+^or Cu^2+^. For sites that AllMetal3D_loc_ predicts which are not close to any experimental metal site, MIB2 is mostly correct in predicting no metal (Figure S4). When taking the best prediction among the top-3 predictions, the results for MIB2 improve slightly with fewer misannotated Mg^2+^, Ca^2+^and Zn^2+^sites without resolving the problem with other transition metal ions.

MIC (***Shub et al., 2024***) is a recent method for identity prediction based on fingerprints of a given protein location using a SVC classifier. We investigated how MIC performs on all real metal sites predicted by AllMetal3D_loc_ excluding false positives, since MIC was not trained to predict them. We used the predicted positions and not the real positions since MIC was designed for validating locations early in the refinement process for cryo EM structure determination (***Shub et al., 2024***) and therefore should be robust with respect to slight deviations in the location.

Two different models are available classifying sites either according to a set of ions/molecules with high frequency of occurrence in the PDB (Na^+^, Mg^2+^, Ca^2+^, Zn^2+^, H_2_O and Cl^−^) or an extended set (including in addition K^+^, Mn^2+^, Fe^3+^, I^−^ and Br^−^) (Figure S5). We found that MIC correctly predicts Ca^2+^ and Zn^2+^ binding sites but performs less well for Mg^2+^. Many real metal binding sites (especially those involving Mg^2+^ and Ca^2+^ ) are also predicted as water. Concerning the performance of the extended model, we find that rarer metal ions such as Mn^2+^ or Fe^3+^ are mainly predicted as Zn^2+^.

### Geometry prediction

We used the FindGeo annotations for each site in the dataset to perform an additional classification task based on the fingerprint learned by AllMetal3D_base_ centered on the location predicted by AllMetal3D_loc_. Both AllMetal3D_identity_ and AllMetal3D_geometry_ were trained jointly by summing both loss functions before backpropagation. We reduced the 50 classes from FindGeo to 6 majority labels (Table S2) by combining the same geometries of ideal and slightly distorted variants as well as those with and without vacancies into single classes. Together with a class for false positive predictions from AllMetal3D_loc_, the model thus predicts 7 classes in total.

Figure 5 shows that the AllMetal3D_geometry_ model differentiates well between tetrahedral and octahedral geometries. However, the model also frequently mispredicts other categories such as irregular, square or pentagonal bipyramid as octahedral. Tetrahedral sites on the other hand have fewer mispredictions. The other category with few examples in the training set including for example linear (29 training examples) or trigonal prism (27 training examples) is seldomly predicted. Only 35% of the wrong predictions from AllMetal3D_loc_ are predicted correctly by AllMetal3D_geometry_ as NoMetal, on the other hand many real metal sites are predicted as `NoMetal`.

**Figure 5.**
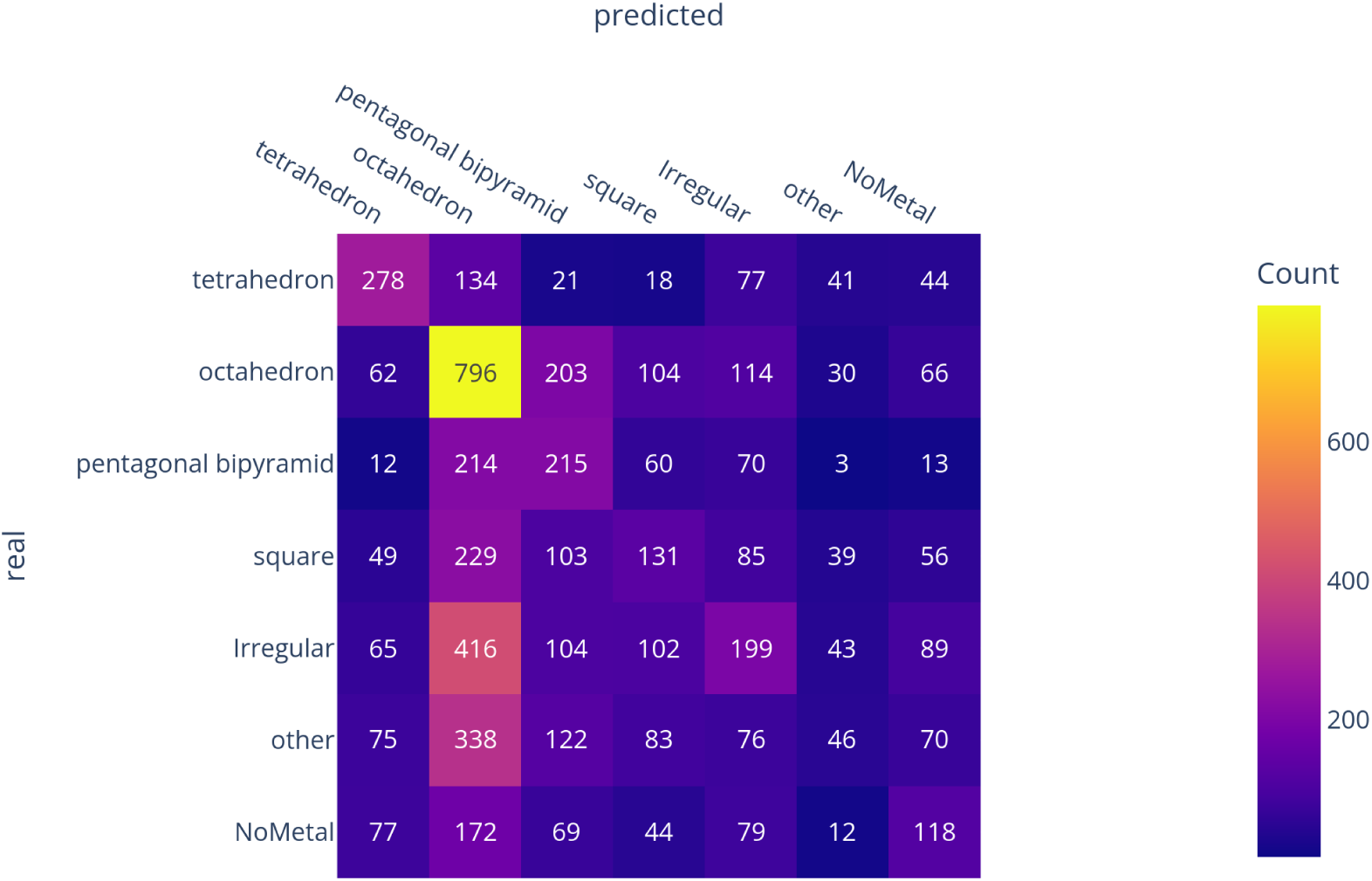
Confusion matrix for geometry prediction on predicted sites from AllMetal3D_loc_.

We also compared AllMetal3D against MetalHawk (***Sgueglia et al., 2024***) on a smaller set of metal sites removing the false positive predictions and non-transition metals for testing since MetalHawk was not trained on these types. We ran MetalHawk both on the predicted locations from AllMetal3D_loc_ as well as on the ground truth sites (Figure S6). Overall AllMetal3D_geometry_ performs much better in differentiating octahedral from tetrahedral sites than MetalHawk. However, similar to what is observed with the larger test set, AllMetal3D_geometry_ frequently predicts other geometries as octahedral. Performance for MetalHawk is marginally improved for tetrahedral geometries when using the real metal locations (Figure S6).

In addition, we ran FindGeo on the AllMetal3D_loc_ location predictions close to true metal ion binding sites and for each real metal site, the closest prediction was used for the FindGeo prediction (Figure S7). While FindGeo accurately identifies most tetrahedral sites given the predicted locations, for all other geometries, predictions are less accurate and the most commonly assigned geometry by FindGeo for the predicted locations is `Irregular`. Octahedral sites are frequently predicted either as square planar or irregular.

### Vacancy prediction

Another important task for applications such as enzyme design is the prediction of sites with vacancies. FindGeo annotates predicted geometries as irregular, fully coordinated or with vacant sites. Based on the fingerprint of AllMetal3D_base_, we also trained AllMetal3D_vacancy_ to predict these geometry labels.

However, we were unable to train such a model due to the data imbalance of irregular and NoMetal sites compared to fully coordinated sites, in fact the trained model always predicts all sites as irregular (Figure S11).

### Inference app

Using the Gradio (***Abid et al., 2019***) library and USCF ChimeraX (***Meng et al., 2023***), we implemented two easy to use applications to run and analyze predictions interactively. In the webapp, users can click on each predicted metal location modelled inside the probability density to view the detailed output for each predicted location. In ChimeraX, users can explore predictions using a table UI widget and keyboard navigation. As an example (shown in Figure 6), for hCA (PDB 2CBA), the physiological metal ion is the sole prediction above p>0.5. The identity classification model correctly predicts the site as Zn^2+^ with non-zinc transition metals ranked second. The geometry model predicts the site as tetrahedral with 30% probability with octahedron ranked second with 19% probability.

**Figure 6.**
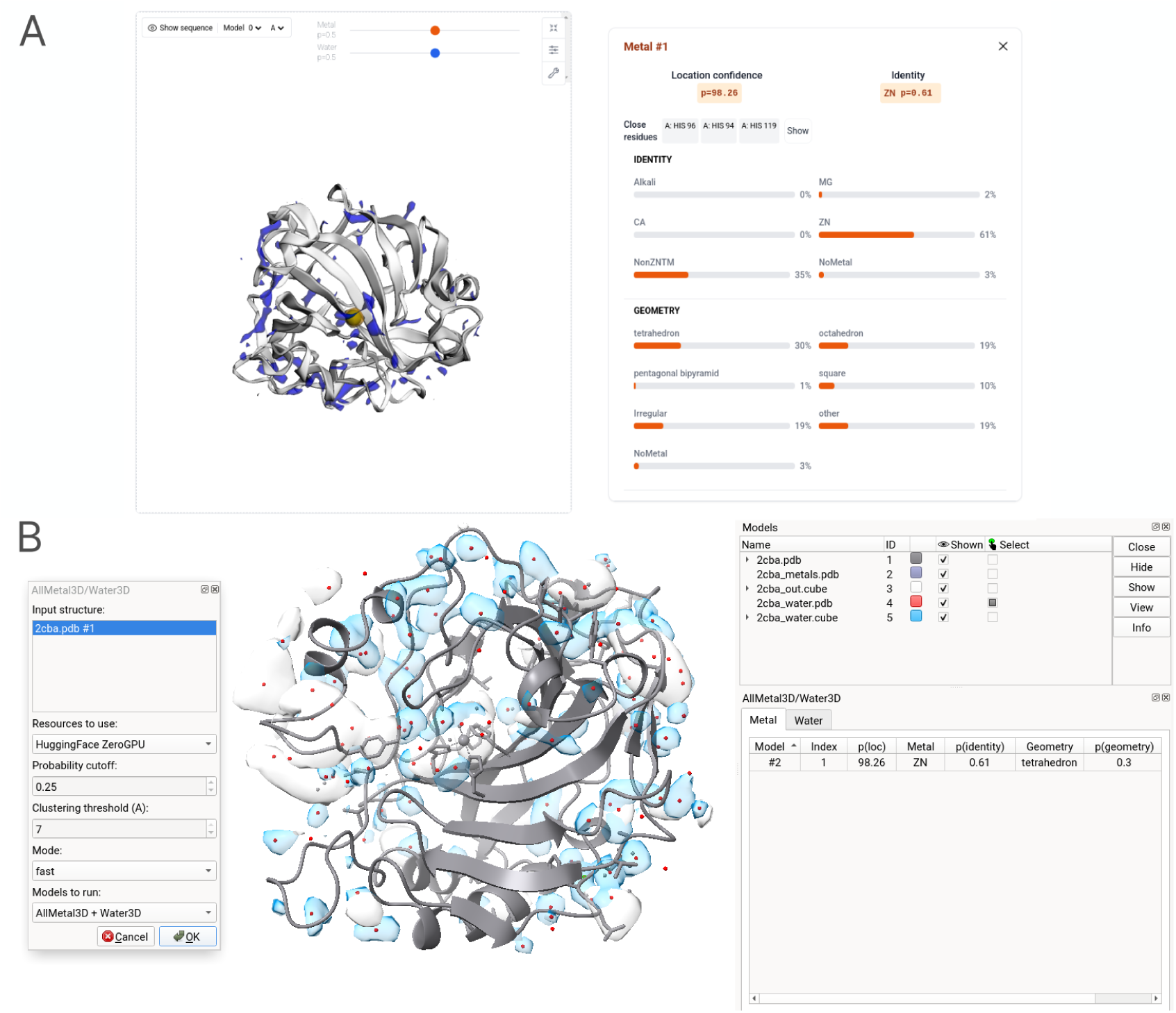
Location, identity and geometry prediction for metals as well as water prediction for carbonic anhydrase using the A Gradio based inference application and the ChimeraX extension B.

### Combined localization and identity model for metals using a unified model

In the original Metal3D paper, we hypothesized that a model for selective location prediction (AllMetal-3D_universal_) could be trained by changing the number of output layers of Metal3D. The two main modifications to Metal3D were the modification of the last layer of the Metal3D model and a change of the training dataset to include all biologically relevant metal ions to create AllMetal3D_universal_. At contrast to the sequential workflow with a specialized location model and classification of the predicted general binding sites we could not train a unified model. Our attempts included different hyperparameter combinations and also using a reduced set of labels for the identity from 12 to 6 labels (data not shown). In contrast to Metal3D, for AllMetal3D_universal_ the size of the output tensor is much larger and sparser during training. All models we trained had consistently decreasing loss. However, all trained models consistently predicted values close to 0 for each point in the input space in inference. This does not yield useful predictions but also fulfills the objective of the binary cross entropy loss because the background signal is much stronger for the larger dataset than for Metal3D trained just on zinc. We also attempted to use a focal loss function to focus on correct predictions against the background but were also unsuccessful.

## Discussion

In this work, we developed a new model for combined location and identity prediction for metal ions in proteins based on the framework of Metal3D (***Dürr et al., 2023***).

### Location prediction

In comparison to Metal3D, AllMetal3D is trained on and is able to predict the location of a wider array of metal ions. The performance for location prediction of Ca^2+^ is now similar to the performance observed for transition metal ions. Performance of K^+^, Na^+^ and Mg^2+^ is improved compared to Metal3D but still not on par with the one for transition metal ions. While for K^+^ this can be expected to a certain extent in view of the relative scarcity of available data, it is surprising that performance for Na^+^ and Mg^2+^ only slightly improves. A possible reason for bad performance on Na^+^ might be that it is often confused with water molecules leading to data quality issues for this metal (***Shub et al., 2024***; ***Zheng et al., 2017***). In fact, ***Shub et al. (2024)*** found that about 20% of sodium is misannotated in the PDB. Na^+^ and K^+^ often use backbone coordination and can adopt a wider array of coordination numbers further influencing data quality. Additionally, Na^+^ with effective ionic radius of 1.02 Å and K^+^ with effective ionic radius of 1.38 Å are larger than most transition metals (e.g. Zn^2+^ 0.74 Å or Cu^2+^ 0.73 Å) (***Shannon, 1976***). This size difference could be a possible reason why there is a performance difference between Na^+^ and K^+^ and the transition metals. We did not adapt the filter sizes that the network uses and applied a constant filter size of 1.5 Å except for the long range filter of 8 Å. It might be that Na^+^ and K^+^ require different filter sizes or that the capacity of the location prediction model needs to be further increased so that it can learn good kernels for the two different sizes of metal ions. The difference in performance of Mg^2+^ and Ca^2+^ is more peculiar given that Mg^2+^ has a lot of training examples available and tighter coordination requirements strictly preferring 6 ligands (***Yang, 2011***). A possible reason why the performance for Mg^2+^ is lacking behind is that DNA containing proteins were excluded from the dataset which likely excludes many of the well defined Mg^2+^ binding sites. Mg^2+^ uses more general binding motifs with aspartates and glutamates than the transition metals using mainly histidines and cysteines. Since these residues are more frequent in proteins (***Vacic et al., 2007***), it might be harder to differentiate metal binding aspartates and glutamates from non-metal binding ones. Additionally, ***Bazayeva et al. (2024)*** found that in the PDB there are nearly the double amount of Mg^2+^ binding sites with low CN of 1, 2 or 3 than there are for Ca^2+^ binding sites. To improve the predictive performance on Mg^2+^ and Na^+^ the training examples should likely be more carefully chosen instead of sampling all sites available in the structures we selected for training.

### Adapting Metal3D for identity prediction

For Metal3D, we speculated that a model could be trained to predict individual metal channels given that 3DCNNs were shown to work individually on location prediction (Metal3D) and identity prediction (MetalSiteHunter). However, we were unable to train such a universal model, likely because the target output was too sparse with most environments containing just one metal in one out of 7 or 12 channels. This heavily incentivized the model to predict the background label of 0 in order to fullfill the objective of the binary cross entropy loss function. It is likely that a model that performs semantic segmentation to predict a per identity probability density for each metal can only be trained by modifying the architecture (e.g.using a UNet) or by modifying the loss function. Examples used in the computer vision literature are focal losses to upweight correct identification (***Bokhorst et al., 2023***; ***Amid et al., 2019***) or other loss functions such as the soft-Dice loss function (***Sudre et al., 2017***). While we have attempted to use a focal loss function further exploration is necessary. It also became evident during our analysis that existing models for these tasks such as MIC or MetalSiteHunter also face similar problems for per ion classifications. MetalSiteHunter (***Mohamadi et al., 2022***), which the authors claim is highly selective for Zn^2+^, Fe^3+^, Ca^2+^, Na^+^, and Mg^2+^ is not very selective for Mg^2+^ predicting it as the more populated similar ion Ca^2+^. In our analysis for MetalSiteHunter, the false positive locations from AllMetal3D were often also annotated as metal binding. For the negative examples we constructed using a similar method as the authors of MetalSiteHunter, we found that MetalSiteHunter had a strong bias for predicting them wrongly as metal binding. The code used to extract the pocket locations is not provided by ***Mohamadi et al. (2022)***. They centered the input environments on all C_*α*_ atoms of the residues that form the metal binding site but do not specify how the binding site is exactly defined. They also do not specify which center is used for non-metal pockets. In the absence of this information, we use all residues that have any atom within 2.8 Å of the metal ion binding site, respectively the pocket centers defined by fpocket to define the centers used for MetalSiteHunter predictions. Since this choice of definition should only lead to minor variations compared to the centers used by ***Mohamadi et al. (2022)***, we expected to see a similar performance compared to results the authors report. In contrast, the difference between our and their reported results is pronounced and we hypothesize that MetalSiteHunter is likely overfit to its training set and can detect non-metal binding sites only if they look similar to the non-metal binding sites it has seen during training. The MIC method by ***Shub et al. (2024)*** performs a similar task compared to MetalSiteHunter but does not use a negative channel, which likely also leads to overfitting. Further support to this conjecture is lend by the authors’ observation that the model predicts waters in the vicinity of cations also as cation (***Shub et al., 2024***). MIC cannot properly identify rarer classes such as Fe^3+^ in our testing, which well justifies why the authors of CheckMyBlob (***Kowiel et al., 2018***), PinMyMetal(***Zheng et al., 2024***) and also our AllMetal3D_identity_ model only predict merged classes of chemically similar ions. Indeed, the merging of classes makes sense in light of the observation we made for Metal3D, which was able to predict other transition metal ions despite being trained on zinc only (***Dürr et al., 2023***). An additional reason why models should not be trained on selectively predicting exact identities for transition metal ions can be related to the fundamental biochemical properties of metal binding sites where selectivity is often enforced not by the site itself (i.e. information the model has access to). The selectivity of a protein for a specific metal rather is the result of compartmentalization in the cell and the spatial proximity of metal transport proteins when the protein is synthesized that locally increase the concentration of the physiologically relevant ion (***Waldron and Robinson, 2009***). This is information models based on individual structures as input do not have access to. AllMetal3D_identity_ shows that some differences between Zn^2+^ and non-zinc transition metals exist because it can differentiate both classes reasonably well but that there also exists some confusion between these two classes. If multiple different metal ions are present in the buffer, metal binding sites for transition metals will be populated by the thermodynamically most stable metal ion according to the Irving-Williams (***Irving and Williams, 1948***) series. The recent design of an anti-Irving-Williams selective metal binding protein uses a new-to-nature coordination motif also pointing to the fact that such selective binding motifs might not exist in natural proteins (***Choi and Tezcan, 2022***).

MIB2 is the only other method compared to AllMetal3D_loc_ and AllMetal3D_identity_ that can perform blind prediction of metal ion location and identity for transition metals and earth-alkaline metals. However, in our previous work (***Dürr et al., 2023***), we have already shown that MIB2 is inferior to Metal3D for Zn^2+^ ion location prediction. In this work, we also show that MIB2 performs worse than AllMetal3D for selective identity and location prediction for a wider range of metal ions. It is interesting that according to the MIB2 predictions, not the most common ion in the database is preferred (Zn^2+^) but rather often Zn^2+^ is misclassified as Fe^3+^ or Cu^2+^. Taking top-3 predictions also confirmed that the t-score is not directly predictive of metal ion identity.

While for Zn^2+^, we have shown that data quality is not an issue (***Dürr et al., 2023***), for other metal ions this problem might be more pronounced, especially for the task of identity prediction where we could not perform data augmentation as easily as for location prediction where we performed per residue prediction thus allowing us to sample the same metal ion multiple times with different residues centered in the box. For identity prediction on the other hand, the voxelization box is centered on the maximum of the predicted probability density from AllMetal3D_loc_. As a consequence, we could only use rotational data augmentation. The authors of MIC also argue that dataset quality is an issue for classification tasks (***Shub et al., 2024***).

We noted for Metal3D that even low confidence predictions often indicate physically realistic coordination sites that in fact might be populated at high metal load in the buffer. However, to train the identity model, no labels for these ions are available. Therefore we trained the identity model to classify these peaks as false positive (`NoMetal`). Although AllMetal3D_identity_ also receives the probability prediction of AllMetal3D_loc_ as input, overall, it could not faithfully correct these ”wrong” predictions. This further points to a lack of true difference between these predictions and real metal binding sites. These false positives could indeed be real metal ion binding sites that just were not yet resolved in the experimental structure for a variety of reasons such as absence of that metal in the buffer or due to high *β*-factors or processing conditions that lead to the loss of more weakly bound ions.

Overall, AllMetal3D predicts lower maximum probabilities for the different metal ions we analyzed compared to Metal3D and fewer low confidence predictions that could indicate potential low affinity metal binding sites. Both of these observations can likely be linked to the more heterogeneous dataset with multiple ions with different binding modes. The signal to noise ratio in the metal dataset of AllMetal3D is likely decreased compared to the more homogeneous zinc dataset of Metal3D.

### Geometry and vacancy prediction

We hypothesize that the inferior performance for geometry and vacancy prediction can be linked to the training set diversity as well as the slight geometric inaccuracies. While for geometry prediction the model can at least differentiate between tetrahedral or octahedral sites, our attempts at predicting vacancies were entirely unsuccessful. Since the difference between coordination geometries of metal ion binding sites can be quite small (for example between square pyramidal and tetragonal bipyramidal), differentiating between them can be difficult for the higher coordinated geometries (***Andreini et al., 2012***). The lack of rotational equivariance of our models together with the spatial imprecision of the AllMetal3D_loc_ predictions on the order of 0.5 Å further interferes with accurate geometry assignment. In addition, crystal structures especially the ones above 2.5 Å resolution usually also have positional inaccuracies for the metal ion location resulting in difficulties in differentiating similar geometries. It is therefore unclear how good the ground truth FindGeo (***Andreini et al., 2012***) labels are. By running FindGeo on the predicted locations, we were able to confirm that FindGeo has difficulties predicting the same geometries as for the experimental locations. This shows that slight geometric deviations indeed cause frequent misassignments. This makes it harder for the model to predict the ground truth label and contributes to the lack of performance in classifying rarer geometries.

We also tested the performance of MetalHawk (***Sgueglia et al., 2024***) trained on the CSD dataset, which uses a much simpler neural network than AllMetal3D. In our testing, MetalHawk had little correlation with the FindGeo labels. In agreement, the authors of MetalHawk (***Sgueglia et al., 2024***) also found that when they trained on a dataset created from the PDB, their model did not perform well because of limited data on rarer classes. The worse performance of MetalHawk could however also be rooted in the slight geometric deviations in the predicted locations that we found to influence the FindGeo predictions. MetalHawk probably is very sensitive to these deviations since it uses distance inputs. The disagreement between the FindGeo assignments and MetalHawk on experimental locations further points to deficiencies in the MetalHawk model.

### Cofolding methods

New cofolding methods such as RoseTTAfold-AllAtom RFAA) (***Krishna et al., 2024***) and AlphaFold3 (AF3) (***Abramson et al., 2024***) also support the prediction of ions analyzed in this work. However, a key difference is that in comparison to AllMetal3D or MIB2, they use a known stoichiometry for metal ion binding. The prediction using these cofolding methods is only blind for the location itself. AllMetal3D performs truly blind predictions given only the structure and not the identity or number of bound metal ions. A detailed comparison of the performance of RFAA/AF3 with respect to AllMetal3D was performed in ***Dürr and Rothlisberger (2024)***.

## Conclusion

AllMetal3D is a pipeline composed of a retrained Metal3D model to predict the location of a wide array of metal ions and a classification network that assigns the predicted probability density peaks to specific metal identities. AllMetal3D predicts metals of several classes such as Zn^2+^, Ca^2+^ or non-zinc transition metal to further aid in their identification. Instead of predicting classes of all individual metal ions, which we found to be prone to overfitting when comparing with other methods, we predict grouped classes of chemically similar metal ions. Both location and identity prediction come with a confidence metric with the identity model being able to predict possible false positive location predictions that can be manually verified. AllMetal3D outperforms both existing methods that predict the location of metal ions as well as models that classify metal ion sites. We expect AllMetal3D to be broadly useful in the context of structure prediction, validation of experimental structures or for the design of novel metalloproteins.

## Methods

In this work, we developed a family of models called AllMetal3D for location prediction (AllMetal3D_loc_) and for various classification tasks (e.g. AllMetal3D_identity_ and AllMetal3D_geometry_). These classification models use a fingerprint representation of each metal site learned by AllMetal3D_base_.

### Dataset

We constructed the dataset for training and testing in a similar fashion as described for Metal3D (***Dürr et al., 2023***). Briefly, sequence clusters from the PDB were extracted at 30% sequence identity using MMSeqs2-easy cluster (***Steinegger and Söding, 2017***). We used the same training cutoff (5^th^ March 2021) as for Metal3D. The highest resolution structure containing one of the target metals was extracted from each cluster. The list of metal PDB codes included is `MG,CA, NA, K, ZN, FE, FE2, CU, CO, CU1`, and `MN`.

We always used the biological assembly to sample metal ions and used the biological assembly and symmetry adjacent copies to construct the environment for voxelization so that metal sites at crystal contacts between periodic replicas of the system are sampled in the fully coordinated *in crystallo* environment. For each structure, we sample all residues around the metal ion and an equal number of residues at least 12 Å from any ion in the structure to balance the dataset. The same voxelization strategy as for Metal3D was used with the grids centered on the C_*α*_ atom of each residue. Three different versions of the dataset were created. Two where the target is an [`n_metals, 32,32,32`] or [`n_metals_remapped, 32,32,32`] tensor with each channel containing the target probability density for a single metal, respectively categories of similar metals, and the third with size [`1,32,32,32`] with all metals lumped together in a single channel.

For each metal in the training set and test set we ran FindGeo (***Andreini et al., 2012***) in a 32-bit docker container to predict the coordination geometry because at the time no 64-bit executable was available. We remapped the detailed geometries from FindGeo according to Table S2. The output from FindGeo was also used to assign binding sites with vacant coordination sites i.e. where FindGeo predicts additional non-protein ligands to be involved in metal binding.

For AllMetal3D_base_, we did not use C_*α*_-centered voxelization but chose to voxelize an environment centered on the predicted density peak from AllMetal3D_loc_. The density peaks were obtained by taking all structures in the dataset for training of AllMetal3D_loc_ and running inference using all residues as input within 2.8 Å or 4 Å (Na^+^ and K^+^) for AllMetal3D_loc_. Each environment was randomly rotated with three-fold data augmentation. For AllMetal3D_base_ the test set contained 6185 metal sites in 2228 structures and the training set contained 9906 metal sites in 4312 structures.

For evaluation a negative dataset of non-metal proteins was constructed using the non-metalloproteins from the sequence clusters used to construct the dataset of metalbinding proteins. Assignment as non-metalloprotein was performed by checking if a metal was contained in the PDB file, thus structures of metalloproteins resolved experimentally without metal ions would be classified as non-metalbinding. However, all examples we referenced with the Uniprot (***The UniProt Consortium, 2015***) database did not have annotation as metal binding. In total, we sampled 550 structures to create a sufficiently large set even for cases where the various methods might fail processing for some proteins. In fact, the most common mode of failure was that MetalSiteHunter does not process PDB files with multiple molecules such as in biological assemblies where multiple copies of the same domain are deposited as multiple models. For MIB2, a smaller dataset was used because of the heavy rate limiting of the server (1 job per minute with a maximum of 2 concurrent jobs every 10 minutes). The center of each pocket was determined using fpocket (***Le Guilloux et al., 2009***) run in a Docker container with default settings. The center of the largest pocket in the protein was determined by taking the geometrical average of all atoms contacted by the Voronoi vertices of the pocket as determined by fpocket.

### Models

#### AllMetal3D_universal_: Combined localization and identity model

We attempted to only change the dataset and modify the output layers of the Metal3D model so that for each metal in the dataset a separate output channel is predicted. We also attempted to reduce the number of output layers to a smaller set of similar metal classes: `MG, CA, Alkali, ZN, NonZNTM, MN. NonZNTM` are all transition metals except for Zn^2+^ and Mn^2+^. We treated Mn^2+^ separately because it often also binds similar as alkaline-earth ions (***Yang, 2011***). We used the same loss function, model architecture and hyperparameters for training as for Metal3D. This model we refer to as AllMetal3D_universal_.

#### AllMetal3D_loc_: Location prediction with Metal3D framework

After failure to train AllMetal3D_universal_ for combined location and identity prediction, we attempted to train a simpler model called AllMetal3D_loc_ (Figure 1). Here, we used the exact same hyperparameters as for Metal3D predicting one channel of metal density for all aforementioned PDB codes and averaging predictions for individual residues to obtain a global map of metal ion binding per protein in the same way as for Metal3D. The model was trained for 6 epochs.

#### Identity, geometry and vacancy prediction for AllMetal3D_loc_ predictions

Subsequently, we developed a second model with different modules for identity, geometry and vacancy prediction based on a fingerprint learned by the base model. We refer to the common part of this model as AllMetal3D_base_ and the fully connected model using the fingerprint for a specific task as AllMetal3D_identity_, AllMetal3D_geometry_ and AllMetal3D_vacancy_, respectively (Figure 1). The base model extracts the environment around the predicted metal position and condenses the features of the environment into a fingerprint. The small classification models use the fingerprint to predict the identity or geometry of the predicted metal location.

The base model is trained by taking all metal ions in the training set and running inference with AllMetal3D_loc_ on these structures by using the C_*α*_-atoms of residues within 2.8 Å or 4 Å (Na^+^, K^+^) of the metal site as center of the voxelgrids. Similarly, the AllMetal3D_loc_ model is used to generate the test set. Predictions using AllMetal3D_base_ are performed only on correctly predicted metals by AllMetal3D_loc_ as well as false positives (i.e. binding sites that are not close to a metal). Each density peak by AllMetal3D_loc_ is labeled with the closest metal ion in the experimental structure. Any density peak not within 5 Å of a experimental metal is labeled as `NoMetal`. All false negative metals (i.e. metals in the experimental structures that are not predicted by AllMetal3D_loc_) are discarded. The final environment for the subsequent processing by AllMetal3D_base_ was voxelized using the maximum in the predicted probability density maps as center and the same input channels as for AllMetal3D_loc_.

AllMetal3D_base_ is composed of a convolution block composed of 5 convolution layers with 8, 60, 100, 80, 30 and 20 channels. After the second channel max-pooling is applied with filter size 2. Between each convolution layer, the leaky ReLU non-linearity was used with negative slope of 0.2 (***Maas et al., 2013***). The features extracted in the convolution part are then concatenated into a fingerprint representation of size 1280. To the fingerprint we added the probability predicted by AllMetal3D_loc_. AllMetal3D_identity_, AllMetal3D_geometry_ and AllMetal3D_vacancy_ are all fully connected networks composed of three layers with sizes 250, 100 and the length of the classes to predict. The leaky ReLU non-linearity was used with the same negative slope of 0.1. We used a stepped learning rate with *γ* = 0.9 and the Adam (***Kingma and Ba, 2017***) optimizer with a default learning rate of 0.001. A batch size of 512 was used. The model was trained for 300 epochs.

The parameters for AllMetal3D_identity_, AllMetal3D_geometry_ and AllMetal3D_base_, which are listed above were obtained using hyperparameter tuning among the parameters reported in Table 1.

**Table 1.**
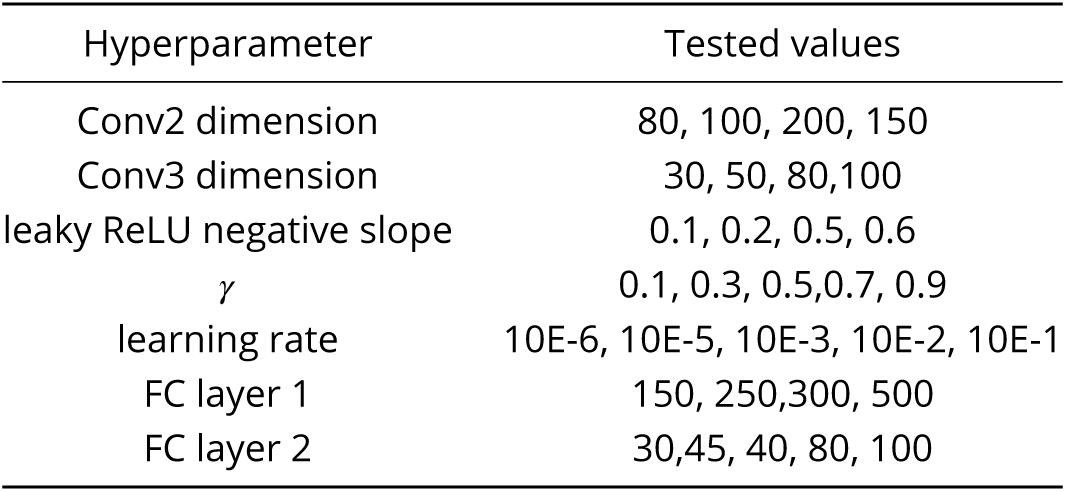
Hyperparameters for AllMetal3D_identity_.

| Hyperparameter | Tested values |
| --- | --- |
| Conv2 dimension | 80, 100, 200, 150 |
| Conv3 dimension | 30, 50, 80, 100 |
| leaky ReLU negative slope | 0.1, 0.2, 0.5, 0.6 |
| $\gamma$ | 0.1, 0.3, 0.5, 0.7, 0.9 |
| learning rate | 10E-6, 10E-5, 10E-3, 10E-2, 10E-1 |
| FC layer 1 | 150, 250, 300, 500 |
| FC layer 2 | 30, 45, 40, 80, 100 |

### Evaluation

The same evaluation metrics to calculate correct (true positive) and wrong (false positive) predictions as well as not found sites (false negatives) are used as for Metal3D (***Dürr et al., 2023***).

#### AllMetal3D

Inference using the trained models is always run in PyTorch eval mode. We always run predictions on all residues in the protein for inference. Two faster modes are also available. With blocked sampling of residues each C_*α*_ atom is included in at least one of the sampled environments to cover the whole protein. It is also possible to run on a given residue and C_*α*_ atoms in a given radius.

#### MetalSiteHunter

We used the publicly available webserver^1^ (no longer available as of February 25) to run predictions. Since no API was available, we employed the clicknium library to automate submitting protein structures to the server. For each structure, we used tampermonkey to insert custom javascript into the webpage to set the center of the analyzed box to either the geometric center of all atoms lining a given pocket (non-metalloprotein dataset) or to the center of all C_*α*_-atoms of the residues directly coordinating the metal using 2.8 Å respectively 4 Å for Na^+^ and K^+^ as the cutoff. ***Mohamadi et al. (2022)*** did not exactly specify the criteria they used to determine the metal binding residues. The probabilities and labels were extracted from the HTML output.

#### MIB2

MIB2 (***Lu et al., 2022***) is only available as webserver^2^. We submitted predictions manually with 2 jobs possible per 10 minutes. We stored the job ids and then extracted the results by parsing the HTML of the output. The following PDB codes were used for the input: CA, CU, FE2, MG, MN, ZN, CD, FE, NI, CO, CU1.

#### MetalHawk

The released version of MetalHawk (***Sgueglia et al., 2024***) only processes a single metal ion in a structure. We therefore used a customized version the authors made available to us that can process all transition metals in a given structure. We ran MetalHawk on all real metal sites included in the test set and on all AllMetal3D_loc_ predictions on those structures using the FindGeo labels as the ground truth.

#### MIC

The public code for MIC (***Shub et al., 2024***) was downloaded from GitHub (version 19^th^ March 2024). Predictions were run on the predicted locations given that MIC was designed for identifying water and ions early in the refinement process wherefore it should be tolerant to slight deviations between the experimental locations and the predicted locations. The default options for MIC were used.

## Data Availability

The code is available under https://github.com/lcbc-epfl/allmetal3d and pip installable using pip install allmetal3d. The ChimeraX extension can be installed from the ChimeraX Toolshed and its code is available under https://github.com/lcbc-epfl/allmetal3d-chimerax-plugin. Data and Code are archived under DOI https://doi.org/10.5281/zenodo.14809847. Documentation is available under https://lcbc-epfl.github.io/allmetal3d/.

## Acknowledgments

U.R. acknowledges funding from the Swiss National Science Foundation under grant No. 200020−219440 and computational resources from the Swiss National Computing Center CSCS.

## Appendix Location model

**Appendix 0—figure S1.**
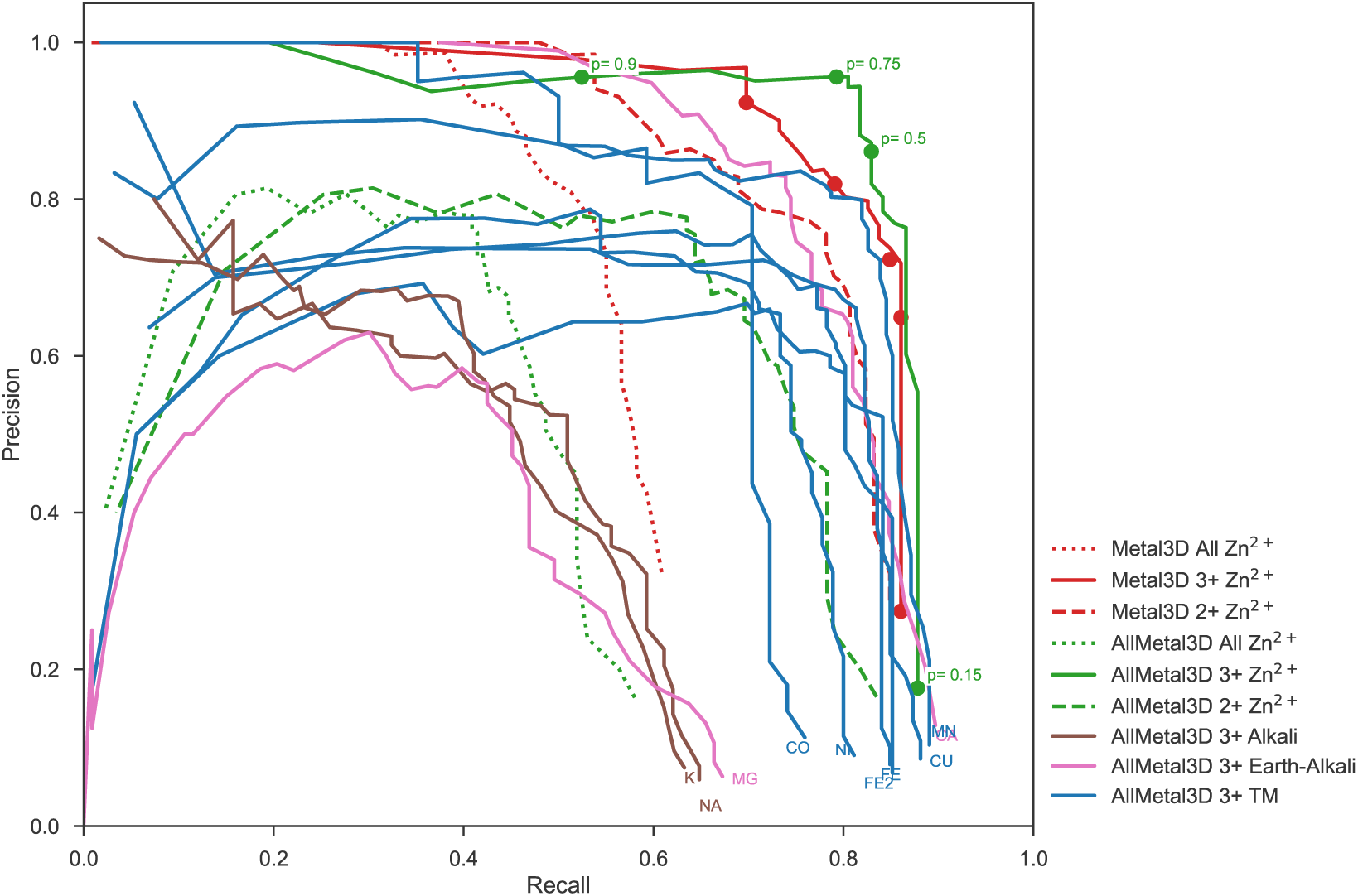
Full precision and recall for metal binding sites for Metal3D and AllMetal3D_loc_ for sets of physiological sites with 3+ unique coordinating residues, metals with 2+ residues and all metal binding sites.

**Appendix 0—figure S2.**
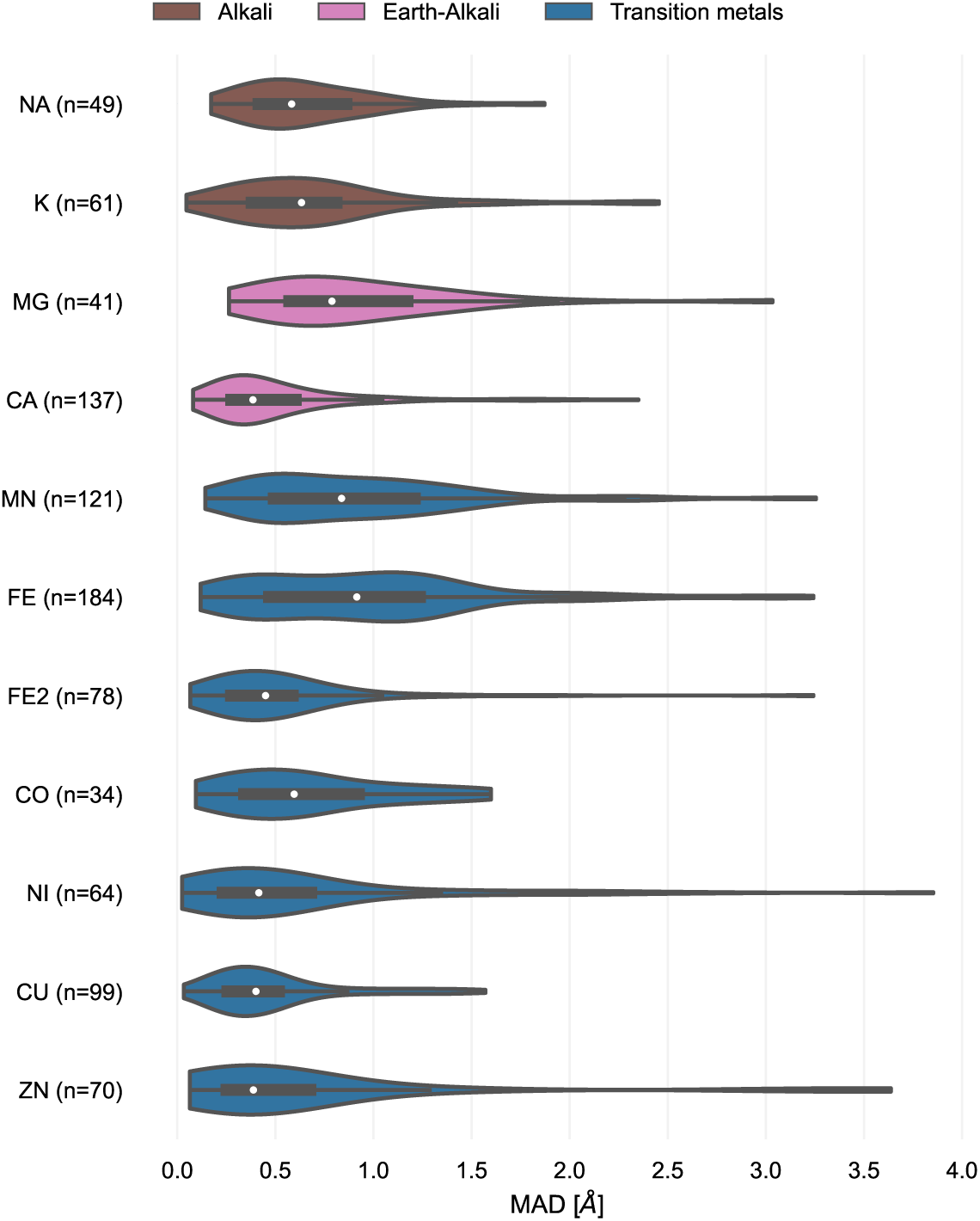
Mean absolute deviation for location prediction for physiological metal sites with 3+ unique protein ligands for AllMetal3Dloc *p* = 0.65.

**Appendix 0—figure S3.**
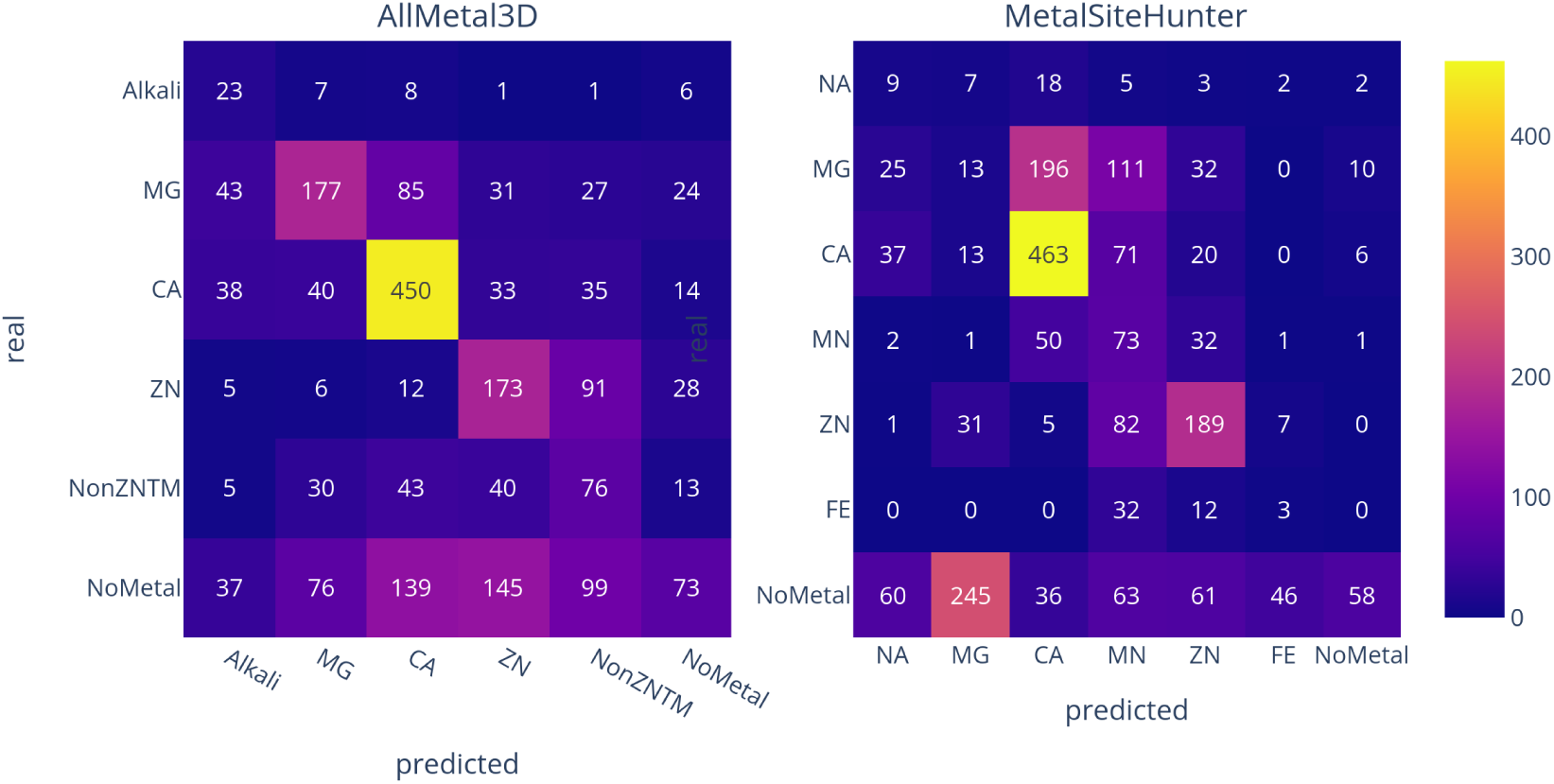
Comparison of MetalSiteHunter and AllMetal3D_identity_ on the same set of 2960 site predictions by AllMetal3D_loc_.

**Appendix 0—figure S4.**
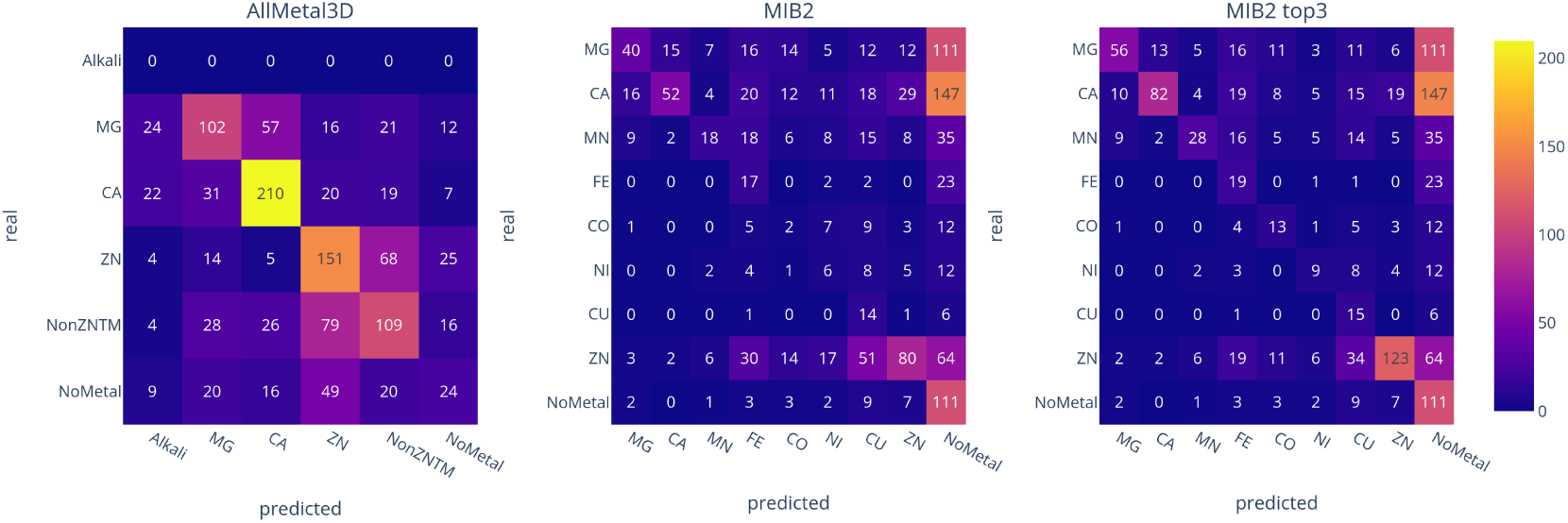
Comparison of AllMetal3D_identity_, MIB2, MIB2 top3 on the same set of 1520 site predictions by AllMetal3D_loc_.

**Appendix 0—figure S5.**
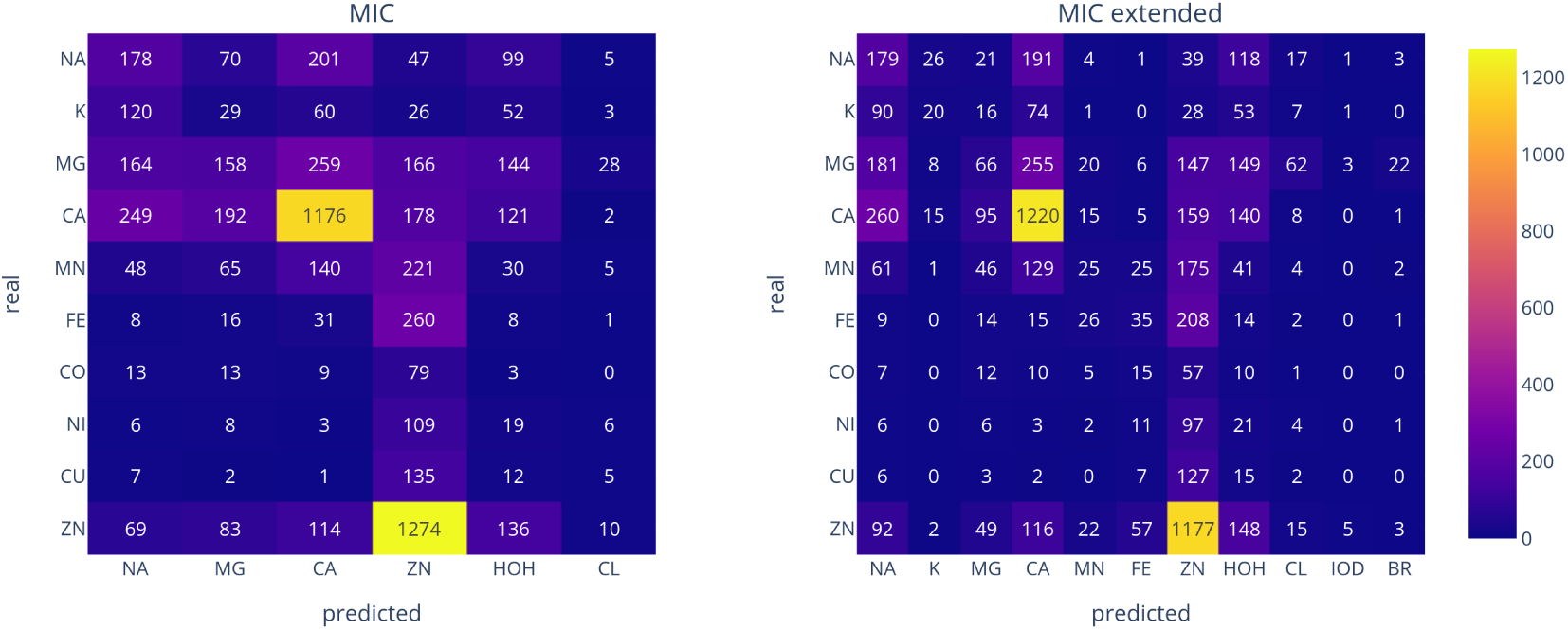
MIC predictions on metal sites present in the AllMetal3D test set excluding false positive predictions by AllMetal3D_loc_.

**Appendix 0—figure S6.**
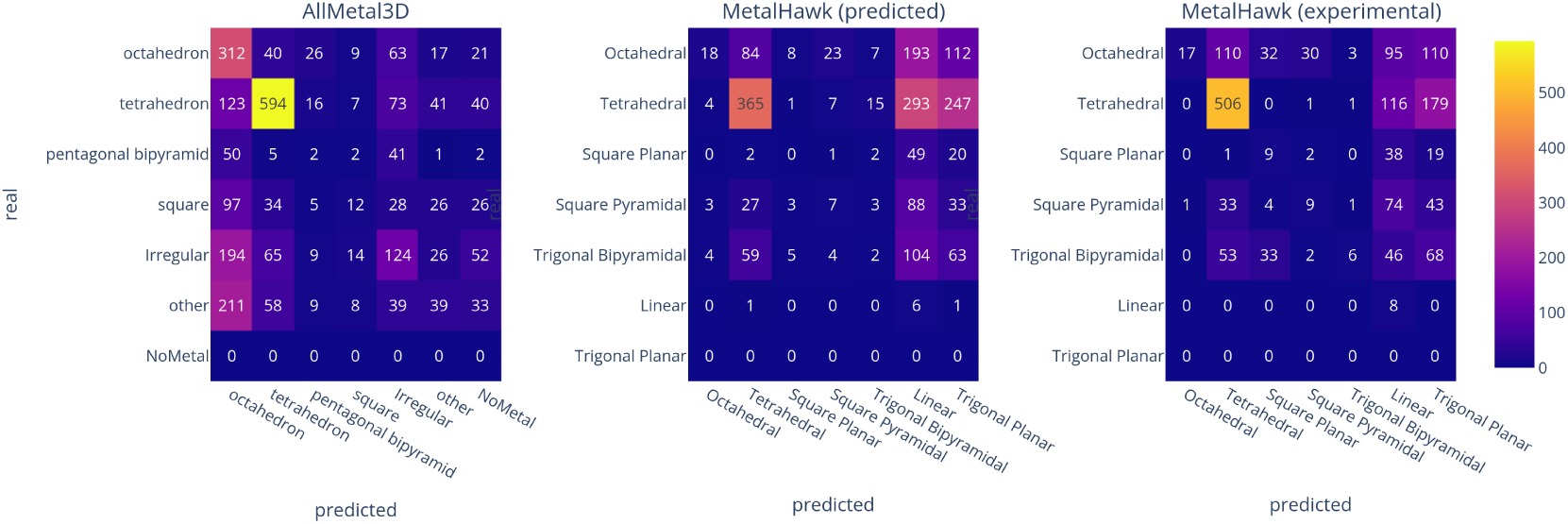
Confusion matrix for geometry prediction from AllMetal3D_identity_ and MetalHawk on predicted locations for real metal binding sites in the test data set from AllMetal3D_loc_ and on ground truth experimental metal locations

**Appendix 0—figure S7.**
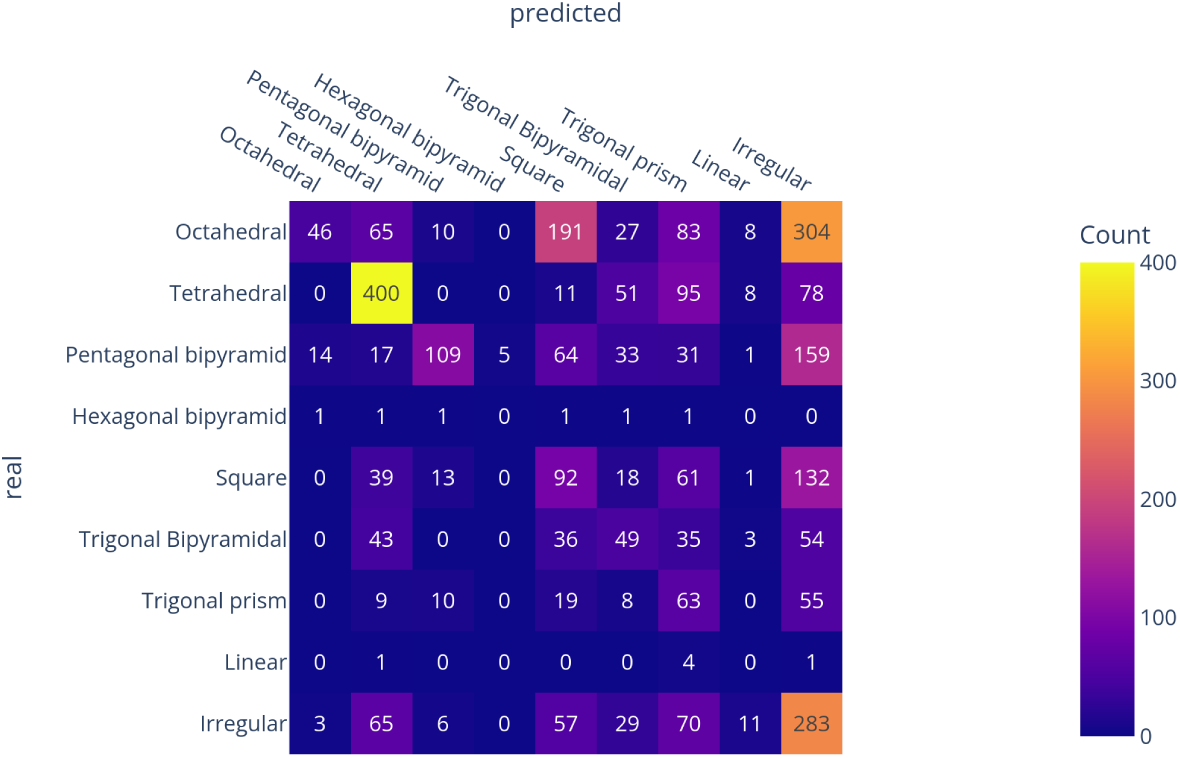
Predictions of FindGeo on locations predicted by AllMetal3D_loc_.

**Appendix 0—figure S8.**
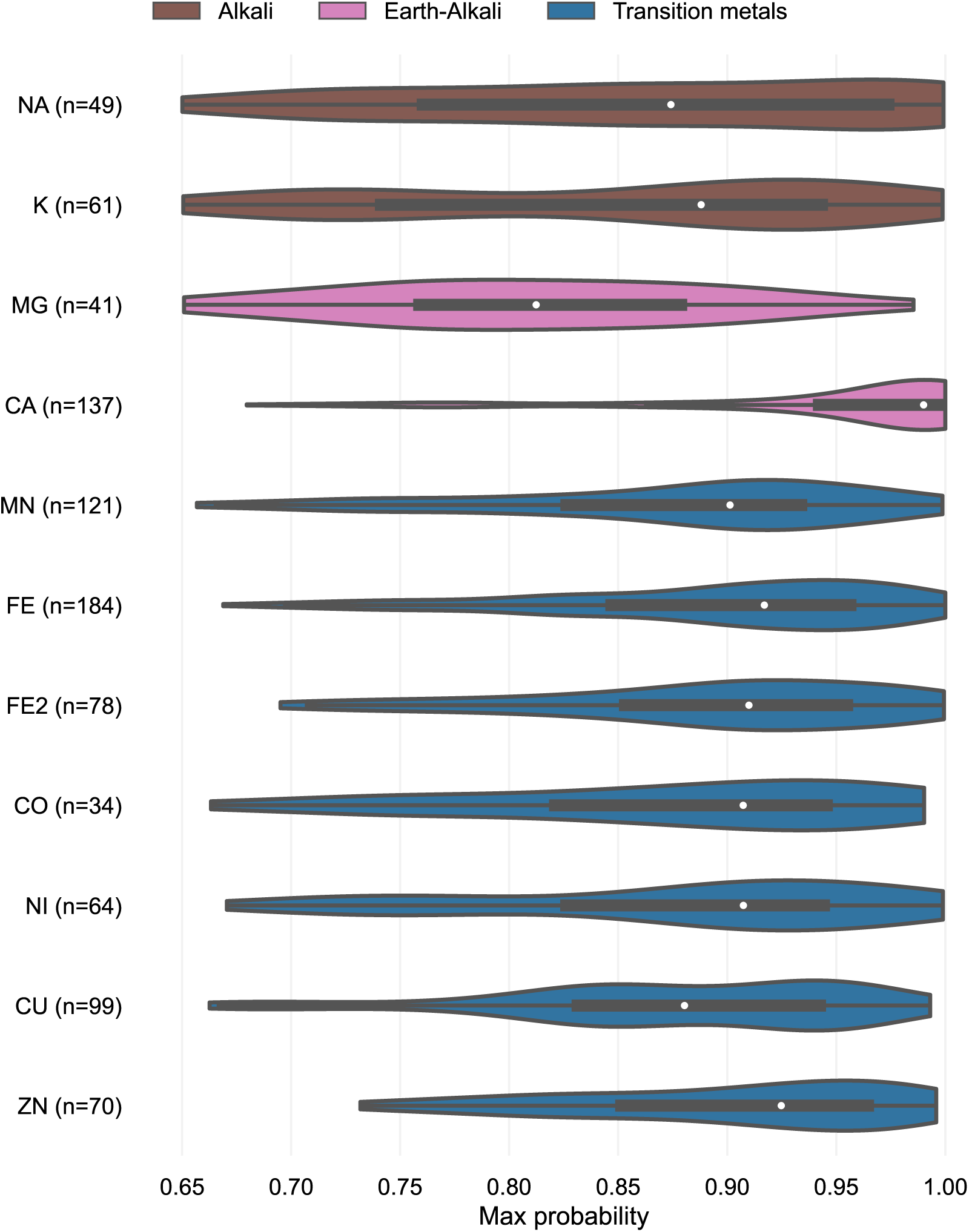
Maximum probability for predicted locations for various metals for AllMetal3D_loc_ p=0.65

**Appendix 0—figure S9.**
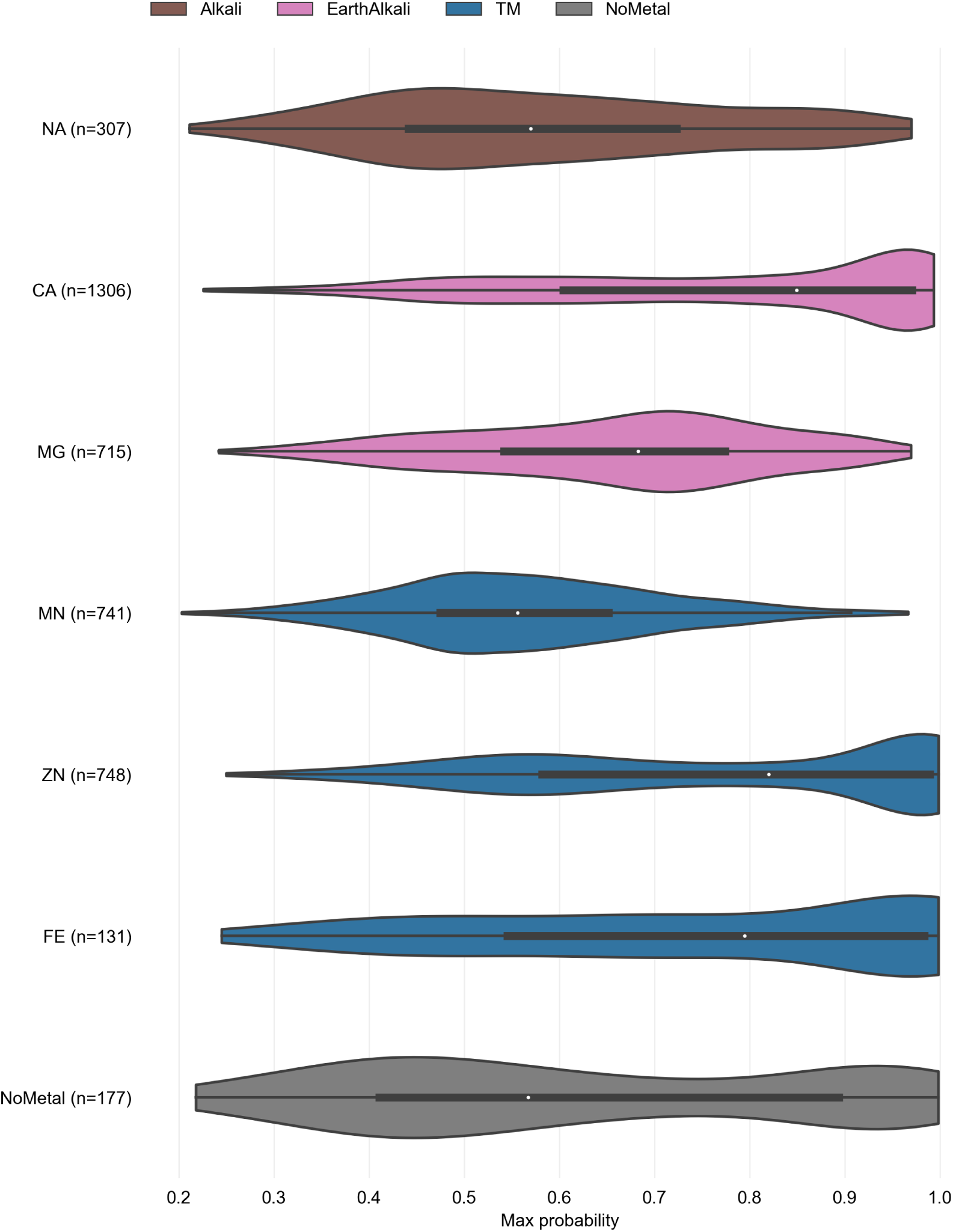
Probability distribution MetalSiteHunter with median indicated by white dot.

**Appendix 0—figure S10.**
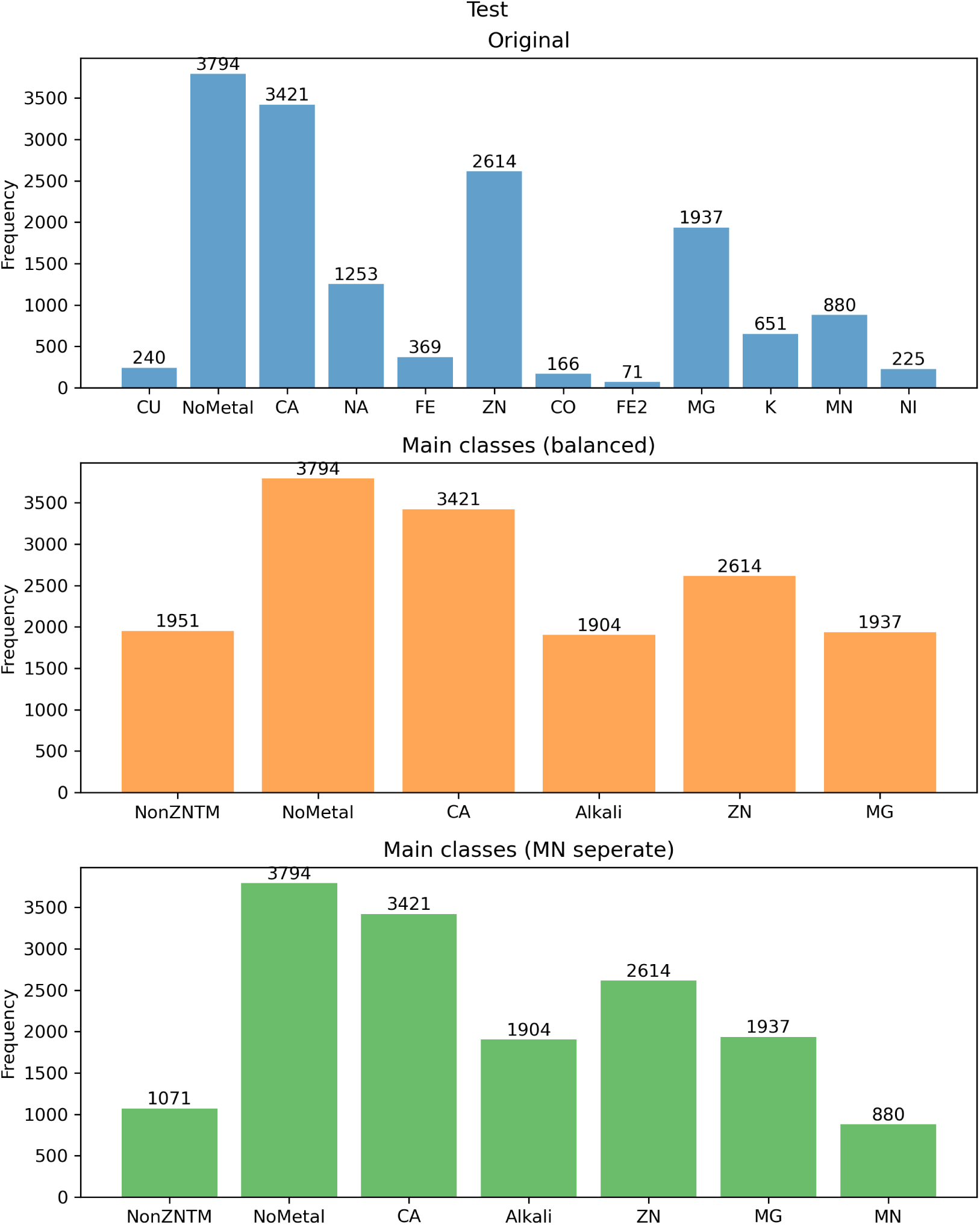
Class distribution in test set of AllMetal3D for original dataset, balanced dataset and separate manganese.

**Appendix 0—figure S11.**
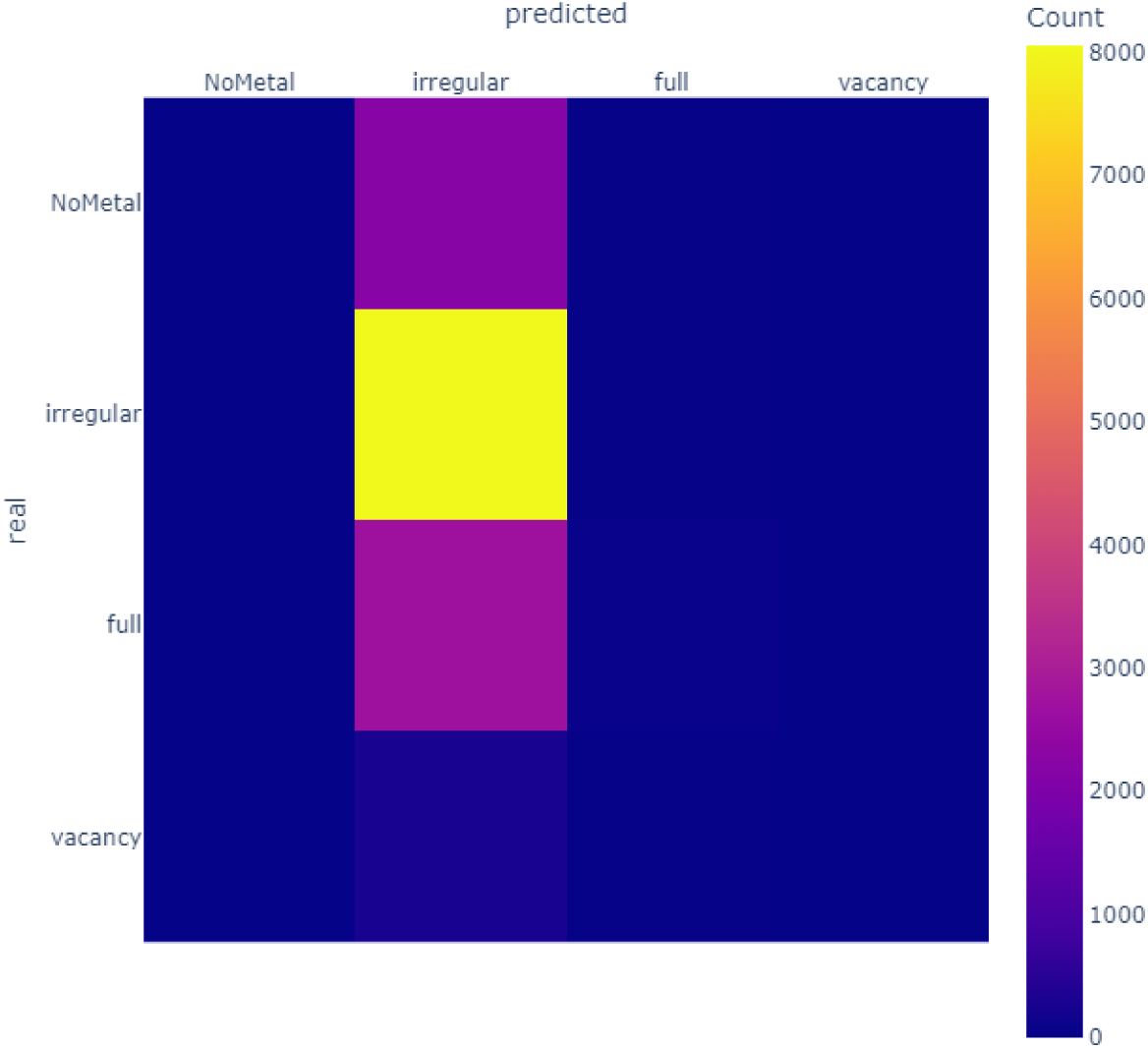
Confusion matrix for vacancy prediction from AllMetal3D_*vacancy*_,

**Appendix 0—table S2.**
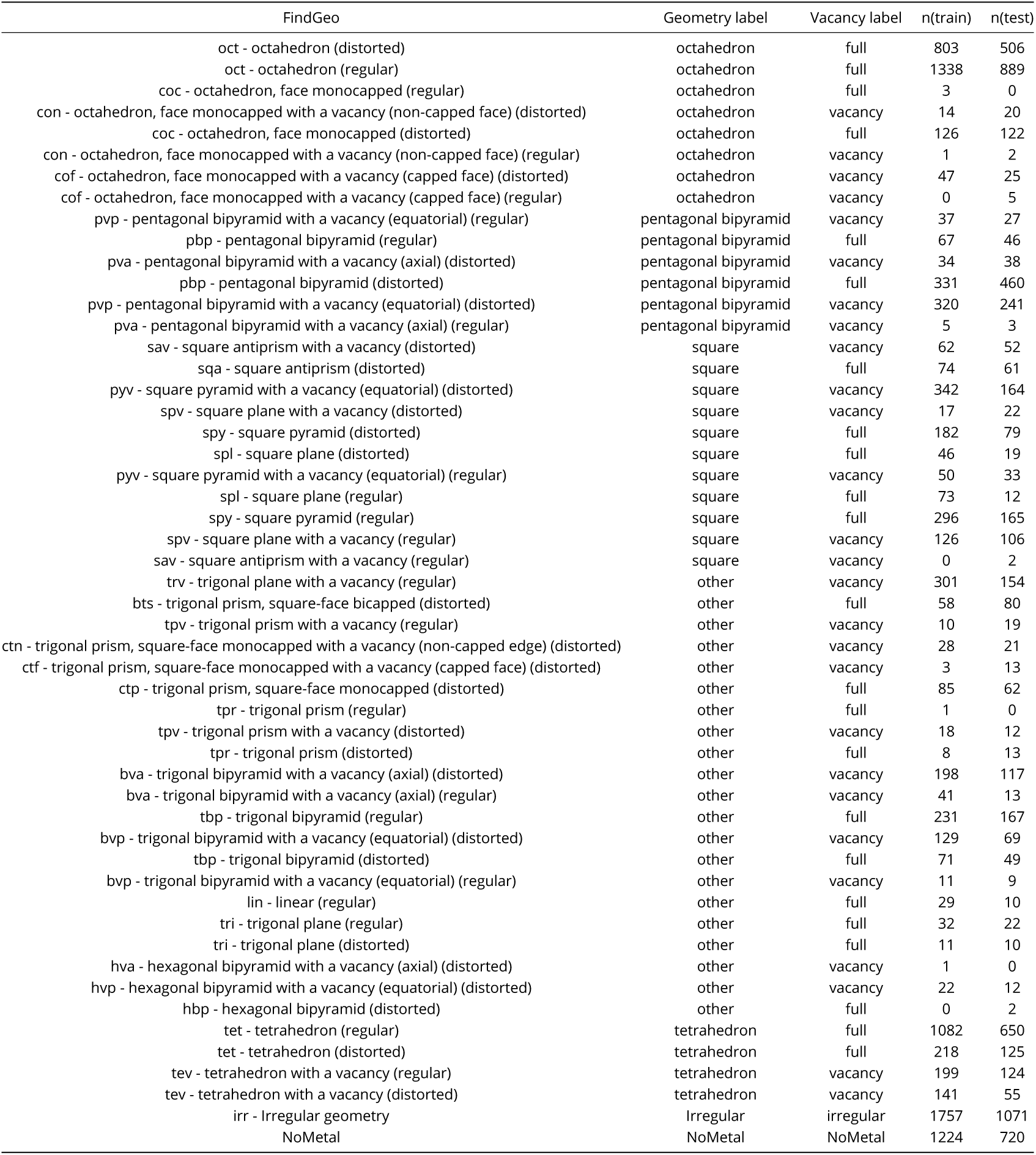
Overview of FindGeo classes present in the dataset for AllMetal3D_*geometry*_ and AllMetal3D_*vacancy*_.

| FindGeo | Geometry label | Vacancy label | n(train) | n(test) |
| --- | --- | --- | --- | --- |
| oct - octahedron (distorted) | octahedron | full | 803 | 506 |
| oct - octahedron (regular) | octahedron | full | 1338 | 889 |
| coc - octahedron, face monocapped (regular) | octahedron | full | 3 | 0 |
| con - octahedron, face monocapped with a vacancy (non-capped face) (distorted) | octahedron | vacancy | 14 | 20 |
| coc - octahedron, face monocapped (distorted) | octahedron | full | 126 | 122 |
| con - octahedron, face monocapped with a vacancy (non-capped face) (regular) | octahedron | vacancy | 1 | 2 |
| cof - octahedron, face monocapped with a vacancy (capped face) (distorted) | octahedron | vacancy | 47 | 25 |
| cof - octahedron, face monocapped with a vacancy (capped face) (regular) | octahedron | vacancy | 0 | 5 |
| pvp - pentagonal bipyramid with a vacancy (equatorial) (regular) | pentagonal bipyramid | vacancy | 37 | 27 |
| pbp - pentagonal bipyramid (regular) | pentagonal bipyramid | full | 67 | 46 |
| pva - pentagonal bipyramid with a vacancy (axial) (distorted) | pentagonal bipyramid | vacancy | 34 | 38 |
| pbp - pentagonal bipyramid (distorted) | pentagonal bipyramid | full | 331 | 460 |
| pvp - pentagonal bipyramid with a vacancy (equatorial) (distorted) | pentagonal bipyramid | vacancy | 320 | 241 |
| pva - pentagonal bipyramid with a vacancy (axial) (regular) | pentagonal bipyramid | vacancy | 5 | 3 |
| sav - square antiprism with a vacancy (distorted) | square | vacancy | 62 | 52 |
| sqa - square antiprism (distorted) | square | full | 74 | 61 |
| pyv - square pyramid with a vacancy (equatorial) (distorted) | square | vacancy | 342 | 164 |
| spv - square plane with a vacancy (distorted) | square | vacancy | 17 | 22 |
| spy - square pyramid (distorted) | square | full | 182 | 79 |
| spl - square plane (distorted) | square | full | 46 | 19 |
| pyv - square pyramid with a vacancy (equatorial) (regular) | square | vacancy | 50 | 33 |
| spl - square plane (regular) | square | full | 73 | 12 |
| spy - square pyramid (regular) | square | full | 296 | 165 |
| spv - square plane with a vacancy (regular) | square | vacancy | 126 | 106 |
| sav - square antiprism with a vacancy (regular) | square | vacancy | 0 | 2 |
| trv - trigonal plane with a vacancy (regular) | other | vacancy | 301 | 154 |
| bts - trigonal prism, square-face bicapped (distorted) | other | full | 58 | 80 |
| tpv - trigonal prism with a vacancy (regular) | other | vacancy | 10 | 19 |
| ctn - trigonal prism, square-face monocapped with a vacancy (non-capped edge) (distorted) | other | vacancy | 28 | 21 |
| ctf - trigonal prism, square-face monocapped with a vacancy (capped face) (distorted) | other | vacancy | 3 | 13 |
| ctp - trigonal prism, square-face monocapped (distorted) | other | full | 85 | 62 |
| tpr - trigonal prism (regular) | other | full | 1 | 0 |
| tpv - trigonal prism with a vacancy (distorted) | other | vacancy | 18 | 12 |
| tpr - trigonal prism (distorted) | other | full | 8 | 13 |
| bva - trigonal bipyramid with a vacancy (axial) (distorted) | other | vacancy | 198 | 117 |
| bva - trigonal bipyramid with a vacancy (axial) (regular) | other | vacancy | 41 | 13 |
| tbp - trigonal bipyramid (regular) | other | full | 231 | 167 |
| bvp - trigonal bipyramid with a vacancy (equatorial) (distorted) | other | vacancy | 129 | 69 |
| tbp - trigonal bipyramid (distorted) | other | full | 71 | 49 |
| bvp - trigonal bipyramid with a vacancy (equatorial) (regular) | other | vacancy | 11 | 9 |
| lin - linear (regular) | other | full | 29 | 10 |
| tri - trigonal plane (regular) | other | full | 32 | 22 |
| tri - trigonal plane (distorted) | other | full | 11 | 10 |
| hva - hexagonal bipyramid with a vacancy (axial) (distorted) | other | vacancy | 1 | 0 |
| hvp - hexagonal bipyramid with a vacancy (equatorial) (distorted) | other | vacancy | 22 | 12 |
| hbp - hexagonal bipyramid (distorted) | other | full | 0 | 2 |
| tet - tetrahedron (regular) | tetrahedron | full | 1082 | 650 |
| tet - tetrahedron (distorted) | tetrahedron | full | 218 | 125 |
| tev - tetrahedron with a vacancy (regular) | tetrahedron | vacancy | 199 | 124 |
| tev - tetrahedron with a vacancy (distorted) | tetrahedron | vacancy | 141 | 55 |
| irr - Irregular geometry | Irregular | irregular | 1757 | 1071 |
| NoMetal | NoMetal | NoMetal | 1224 | 720 |

**Appendix 0—table S3.** Probabilities for correctly identified metal sites for Metal3D and AllMetal3D in the test set.

|  | AllMetal3D | Metal3D |
| --- | --- | --- |
| CA | 0.95 ±0.08 | 0.92 ±0.07 |
| CO | 0.87 ±0.09 | 0.94 ±0.07 |
| CU | 0.87 ±0.08 | 0.96 ±0.05 |
| FE | 0.89 ±0.08 | 0.95 ±0.05 |
| FE2 | 0.89 ±0.08 | 0.96 ±0.05 |
| K | 0.84 ±0.11 | 0.88 ±0.08 |
| MG | 0.84 ±0.09 | 0.89 ±0.06 |
| MN | 0.88 ±0.09 | 0.94 ±0.06 |
| NA | 0.87 ±0.10 | 0.81 ±0.06 |
| NI | 0.89 ±0.08 | 0.97 ±0.04 |
| ZN | 0.90 ±0.07 | 0.97 ±0.05 |

## Footnotes

1 https://mohamad-lab.ai/metalsitehunter/

2 http://combio.life.nctu.edu.tw/MIB2

